# Coordinated Evolutionary Rates in Oxidative Phosphorylation Complexes of Papilionoid Legumes: Cytonuclear Coevolution and Relaxed Selection

**DOI:** 10.64898/2026.05.22.727288

**Authors:** Lydia G. Tressel, Justin C. Havird, In-Su Choi, Tracey A. Ruhlman, Domingos Cardoso, Martin F. Wojciechowski, Robert K. Jansen

## Abstract

Across eukaryotes, mitochondrial (mt) and nuclear genomes coordinate the expression and interaction of gene products to maintain cellular functions. While mitonuclear coevolution has been widely explored in animals, it remains understudied in plants, despite their utility as model systems due to relatively slow mitochondrial evolutionary rates and the presence of plastids. Plants rely on oxidative phosphorylation (OXPHOS) for ATP conversion, which requires cofunctionality and likely coevolution of mitochondrial and nuclear gene products. Here, we investigated evolutionary rate covariation (ERC) between mitochondrial- and nuclear-encoded OXPHOS genes in papilionoid legumes, where plastid-nuclear coevolution and an inversion in plastid DNA have been documented previously. Using 50 legume species spanning 15 papilionoid clades, we estimated evolutionary rates for five gene sets: mt-encoded OXPHOS genes, nuclear-encoded mitochondrial-targeted (N-mt) OXPHOS genes, and three control nuclear gene sets that lack mitochondrial interactions (glycolysis, cell cycle, and cytosolic ribosomal genes). Both mt and N-mt OXPHOS genes exhibited significantly elevated nonsynonymous (*d_N_*) and synonymous substitution rates (*d_S_*) in the 50-kb inversion clade relative to other legumes, suggesting accelerated mitochondrial substitution rates. Moreover, elevated *d_N_*/*d_S_* ratios in mt and N-mt OXPHOS genes in this clade were driven by relaxed purifying, not intensified positive selection. ERCs were highest for OXPHOS complexes and genes with physical mitonuclear interactions, as predicted under mitonuclear coevolution. We discuss how these results compare to other cases of cytonuclear coevolution in plants, including plastid-nuclear coevolution in papilionoids, and why dual-targeted, nuclear-encoded genes that repair mt and plastid DNA may underly patterns of molecular evolution in both organelles.

**Significance Statement:** Mitonuclear interactions are essential for cellular energy production, yet the evolutionary dynamics of these interactions remain poorly understood in plants. This study highlights papilionoid legumes as an important system for understanding how coordinated evolution between mitochondrial and nuclear genes shapes plant genomes. By identifying signatures of mitonuclear coevolution in the 50-kb inversion clade, this work demonstrates how shifts in selective pressures on mitochondrial processes can influence nuclear gene evolution. These findings advance our understanding of cytonuclear coordination in plants and provide a foundation for future studies exploring how interactions among genomic compartments contribute to plant evolution, adaptation, and resilience in agriculturally important lineages.

## Introduction

In mitochondria, the synthesis and functionality of protein complexes require tightly coordinated interactions between nuclear-encoded and mitochondrial-encoded proteins. While the mitochondrial genome (mitogenome) retains a small subset of essential proteins in nearly all eukaryotes, most proteins are encoded in the nucleus, translated in the cytosol, and imported into mitochondria where they must physically interact with mitochondrial-encoded proteins (Rand et al. 2004; Levin et al. 2014). These mitonuclear interactions are key characteristics of eukaryotes (Gray et al. 1999; Sloan et al. 2018), central to cellular energy production (Hill 2015), and a source of genetic incompatibilities that impact fitness and reproductive isolation (Burton et al. 2013; Sloan et al. 2017; Moran et al. 2024).

Nuclear-encoded proteins must interact seamlessly with mitochondrial-encoded proteins to generate functional multisubunit complexes, particularly within oxidative phosphorylation (OXPHOS) complexes critical for electron transport and ATP production. These complexes serve as models for studying mitonuclear coevolution (i.e., joint molecular evolution), as changes in the coding sequences in one genome may drive adaptive changes in the other genome to maintain compatibility (Weaver et al. 2022). Analyses of substitution rates and evolutionary rate covariation (ERC) provide powerful tools to detect such coevolutionary patterns, offering insights into the molecular interactions that sustain energy production across eukaryotes (Clark et al. 2012; de Juan et al. 2013; Forsythe et al. 2021). In particular, ERC analyses are used to detect correlated rates of evolution (e.g., branch lengths in a phylogeny) between two sets of proteins. Strong correlations are predicted if the two sets of proteins are coevolving (Forsythe et al. 2025), although shared selective pressures on co-functioning genes in similar metabolic pathways could also cause high ERC. Finally, if demographic changes (e.g., in effective population size or generation time) are responsible for evolutionary rate variation, ERC should be high across any two sets of genes, even those that are functionally and physically unrelated.

Recent analyses across angiosperms have revealed the widespread influence of mitonuclear coevolution. Lian et al. (2024) employed association analyses to identify significant linkage disequilibrium between mitochondrial and nuclear genomes across seven angiosperm species, providing support for coordinated evolution across genomic compartments. These associations are enriched in nuclear loci encoding mitochondria-targeted proteins, emphasizing their role in maintaining proteostasis and energy production. Similarly, Asar et al. (unpublished data, 2024) reported correlated evolutionary rates between nuclear- and mitochondrial-encoded genes across major clades of land plants, again supporting mitonuclear coevolution in general. These findings provide a context for examining mitonuclear coevolution in specific plant lineages with unusual genomic features.

Complementing comparisons across broad taxonomic groups, studies in closely related flowering plant lineages have provided insights into the specific dynamics of mitonuclear coevolution. In *Silene* (Caryophyllaceae), dramatically accelerated rates of nucleotide substitutions in mitogenomes of select species were accompanied by accelerated changes in nuclear-encoded, mitochondrial-targeted proteins, particularly within OXPHOS complexes, but not in nuclear genes that lack mitochondrial interactions (Havird et al. 2015, 2017).

Mitochondrial OXPHOS evolutionary rates in some heterotrophic plants are also correlated with evolutionary rates of nuclear-encoded genes in the same complexes (Guo et al. 2024). Several other angiosperm clades have experienced accelerated rates of mitogenome evolution (Parkinson et al. 2005; Bakker et al. 2006; Mower et al. 2007, 2012; Zhu et al. 2014; Skippington et al. 2015, 2017; Sloan 2015; Lee et al. 2023) and selection to maintain functional compatibility with rapidly evolving mitochondrial counterparts may drive adaptive changes in nuclear genes, particularly at protein-protein interaction sites within multisubunit complexes.

Fabaceae (legumes) are the third largest angiosperm family, encompassing six subfamilies, ∼765 genera and over 19,500 species (LPWG 2017). Papilionoideae is the largest and most economically important subfamily with 503 genera and 14,000 species (LPWG 2017). This subfamily includes major crop species such as *Glycine max* (soybean), *Lens culinaris* (lentil), *Pisum sativum* (pea), *Cicer arietinum* (chickpea), and *Medicago sativa* (alfalfa), which play critical roles in global food security and agricultural economies (Lee et al. 2021; Choi 2022; Dutta 2022). The combination of ecological significance, economic importance, and extensive phylogenetic diversity of Papilionoideae provides a unique opportunity to examine how mitonuclear coevolutionary dynamics shape genome evolution across a broad and agriculturally relevant lineage.

Most vital crops within Papilionoideae are members of the 50-kb inversion clade, a lineage defined both by its central role in global food production and by a distinctive genomic hallmark—an inversion in the large single-copy region of the plastid genome (plastome) (Doyle et al. 1996; Cardoso et al. 2012; Lee et al. 2021; Choi et al. 2022). This inversion functions as a molecular synapomorphy for the group and has been widely employed in previous phylogenomic studies to clarify relationships within Papilionoideae, although nuclear markers have also reinforced the monophyly of the 50-kb inversion clade (Cardoso et al. 2012; LPWG 2017; Cai et al. 2025). Beyond phylogenetic utility, this clade stands out for its extraordinary species richness and ecological versatility, attributes likely facilitated by advances in symbiotic nitrogen and secondary metabolism, enabling members to adapt across a wide range of habitats (Choi et al. 2022). Complete mitogenome sequencing of various species in the 50-kb inversion clade have provided insights into gene loss, intracellular gene transfer, horizontal gene transfer and considerable variation in mitogenome size, making them an excellent system for examining mitonuclear coevolution (Alverson et al. 2011; Kazakoff et al. 2012; Chang et al. 2013; Naito et al. 2013; Negruk et al. 2013; Bi et al. 2016; Kovar et al. 2018; Shi et al. 2018; Yu et al. 2018; Choi et al. 2019, 2021).

We investigated mitonuclear coevolution and selection pressures in papilionoid legumes, specifically within the 50-kb inversion clade (Wojciechowski et al. 2004). We address four key questions regarding molecular evolution of oxidative phosphorylation protein genes: (i) Are substitution rates correlated in mitochondrial and nuclear-encoded, mitochondrial-targeted genes (N-mt genes) as predicted under mitonuclear coevolution? (ii) Are rates more strongly correlated within OXPHOS complexes or individual genes with direct, physical mitonuclear interactions, as predicted under mitonuclear coevolution? (iii) In lineages where mitochondrial rates are accelerated, are there signatures of positive selection in N-mt genes, as would be expected if changes in mtDNA are driving selection for complimentary changes in nuclear DNA? (iv) Do mitogenomes and plastomes exhibit similar patterns of evolutionary rates, as would be expected if they are under similar selection pressures or governed by similar mechanisms of genome maintenance? Together, these questions aim to uncover the evolutionary dynamics of mitonuclear interactions in this diverse and agriculturally vital legume clade.

## Results

### In the 50-kb inversion clade, d_N_ and d_S_ are elevated in mitochondrial and N-mt OXPHOS genes but not in other gene sets

A maximum likelihood constraint phylogenetic tree constructed using 59 concatenated plastid protein-coding genes (Table S1) for 48 Papilionoideae species and two outgroups was fully resolved with 100% bootstrap support for all but nine nodes (Figure 1). Key clades, including Papilionoideae and the 50-kb inversion clade, were clearly defined and strongly supported (100% bootstrap value). This topology served as the framework for all subsequent substitution rate and coevolution analyses, given that plastid genes are popular markers for plant phylogenetic analyses (Choi et al. 2022) but were not a direct focus here.

**Figure 1.**
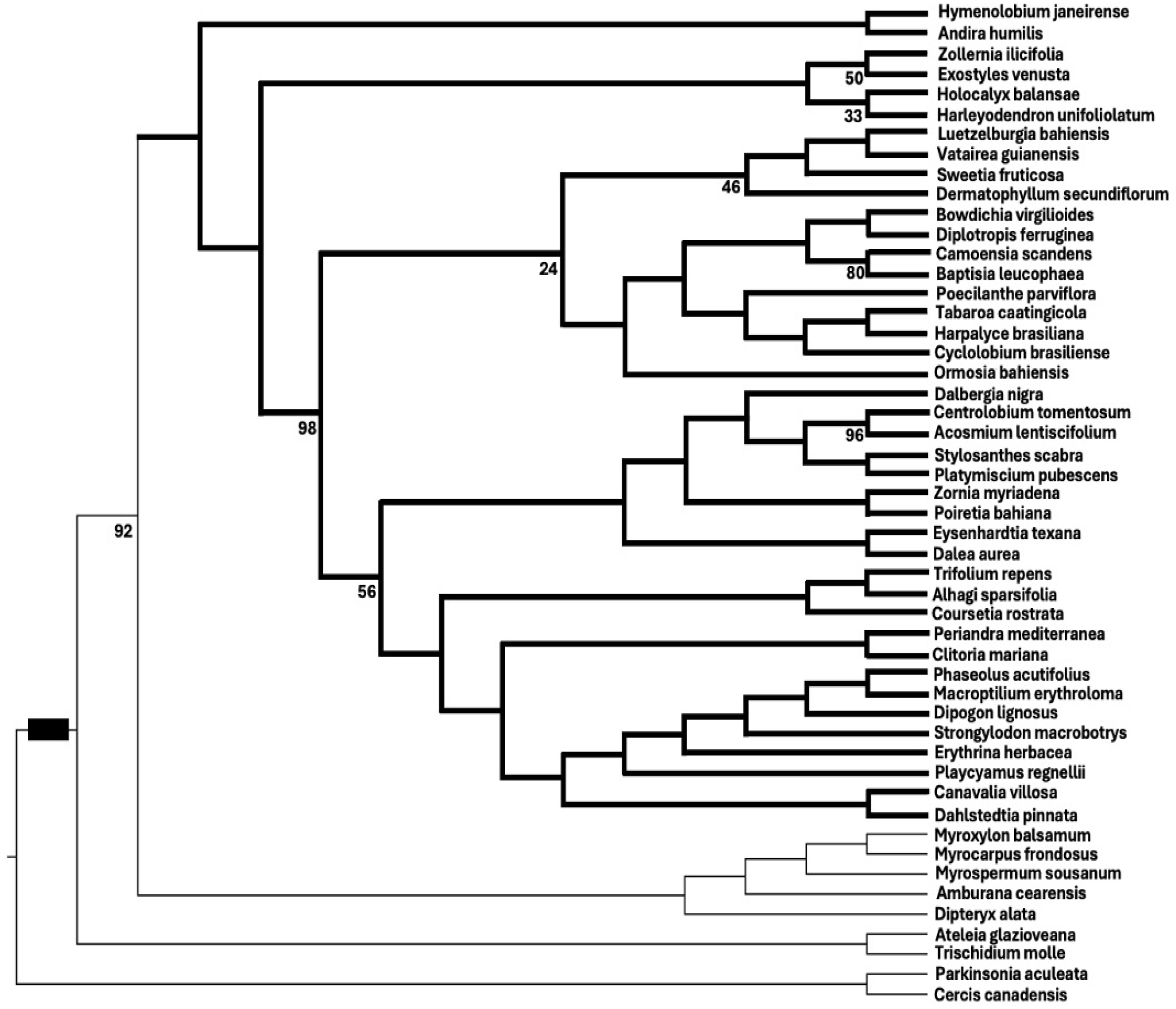
Constraint tree for nucleotide substitution rate estimates and coevolution analyses. The tree topology was based on a maximum likelihood phylogeny generated from 59 concatenated plastid protein coding genes (Table S1). The black rectangle indicates the Papilionoideae clade. The 50-kb inversion clade is specified by the thick lines. All nodes are supported by 100% bootstrap except where indicated.

Nonsynonymous and synonymous substitution rates (*d_N_* and *d_S_*) for all individual genes in mitochondrial and nuclear gene sets were assessed in the 50-kb inversion clade compared to other legumes (model = 2 in PAML). Within the 50-kb inversion clade, both mitochondrial (mt) and nuclear-encoded mitochondrial-targeted (N-mt) OXPHOS genes exhibited significantly higher *d_N_* compared to other legumes using a Wilcoxon rank sum test (*P* < 0.001, Figure 2A). In contrast, three nuclear-encoded control groups of genes with no mitochondrial interactions (glycolysis genes “N-gly”, cell cycle genes “N-cc”, and cytoplasmic ribosomal genes “N-cr”) showed no significant differences in *d_N_* between the 50-kb inversion clade and other legumes (*P* > 0.23 for all; Figure 2A). Similarly, mt and N-mt OXPHOS genes in the 50-kb inversion clade exhibited significantly higher *d_S_* compared to other legumes (*P* < 0.001; Figure 2B). However, *d_S_* were also significantly elevated in the 50-kb inversion clade for the N-gly genes (*P* = 0. 009) and nearly so for the N-cc genes (*P* = 0.065) but was comparable to other legumes for the N-cr genes (*P* = 0.361). We also assessed 20 nuclear-encoded proteins without mitochondrial interactions that were haphazardly chosen from EnsemblPlants (“N-rand” genes, Yates et al. 2022) and there was no significant difference between rates in the 50-kb inversion clade and other legumes for *d_N_*or *d_S_* (*P* > 0.288 for both, Figure 2).

**Figure 2.**
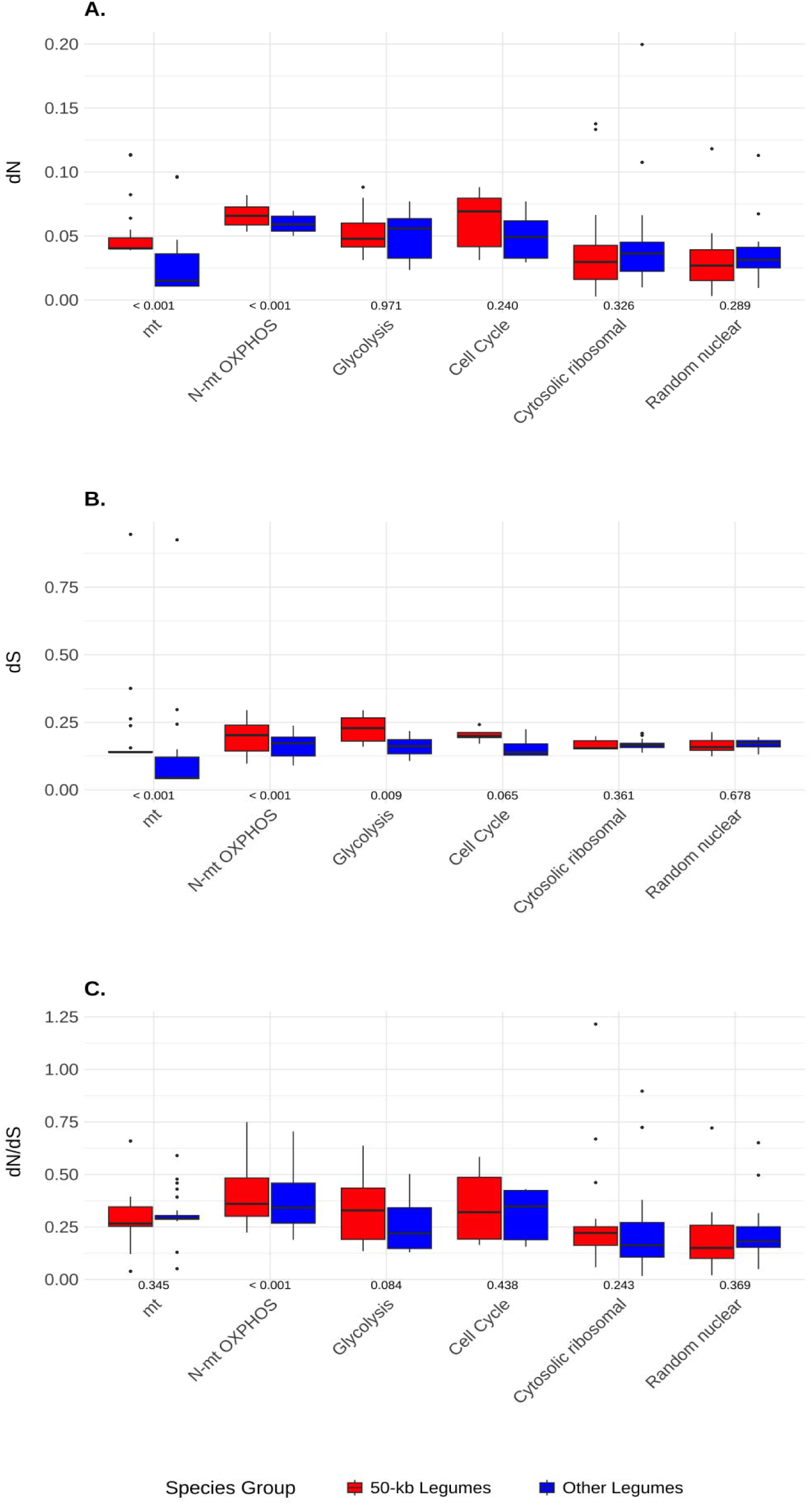
Comparison of nonsynonymous (*d_N_*) and synonymous (*d_S_*) substitution rates and *d_N_/d_S_* ratios in the 50-kb inversion clade vs. other legumes for individual genes in each gene group. Nonsynonymous (A), synonymous (B), and *d_N_/d_S_*ratios (C) were compared across individual genes in different gene groups in the Papilionoideae 50-kb inversion clade vs. other legumes. Gene groups are indicated below the box plots on the x-axis. Outliers are indicated by black dots. *P*-values from Wilcoxon Rank Sum tests after False Discovery Rate correction are below each boxplot comparing the 50-kb inversion clade legumes vs. other legumes in that gene group.

### Nuclear-encoded and mitochondrial-encoded OXPHOS genes show relaxed purifying selection, not positive selection, in the 50-kb inversion clade

*d_N_/d_S_* ratios for individual genes of mt, N-mt OXPHOS, and nuclear gene groups were assessed for signals of positive selection using *d_N_/d_S_* ratios. N-mt OXPHOS genes were the only group of genes that exhibited significantly higher *d_N_/d_S_* in the 50-kb inversion clade compared to other legumes (*P* < 0.001, Wilcoxon rank sum test, Figure 2C), although N-gly genes showed a trend for higher *d_N_/d_S_* in the 50-kb inversion clade (*P* = 0.084). *d_N_/d_S_* ratios were always <1 except for a single outlier cytosolic ribosomal gene, RPS15AE. Likelihood ratio tests (LRTs) were used to further assess whether *d_N_/d_S_* ratios were significantly different between the 50-kb inversion clade and other legumes in our gene sets. Here, a model where a single *d_N_/d_S_* ratio applied to all branches in the phylogeny was compared to a model where *d_N_/d_S_* differed between branches in the 50-kb inversion clade and all other branches. All 60 N-mt OXPHOS and eight of the 17 mt genes exhibited significantly higher *d_N_/d_S_* via likelihood ratio test (*P* < 0.05; Figure 3), whereas the nuclear control genes (N-gly, N-cc, N-cr, and N-rand) showed uniformly high *P*-values consistent with no significant difference in *d_N_/d_S_* ratios between clades. We also performed these analyses on the set of concatenated genes from each gene group, with similar results (Figure 3). Overall, this supports the findings from the Wilcoxon rank sum tests and suggests *d_N_/d_S_* ratios are elevated in the 50-kb inversion clade, but only for N-mt OXPHOS genes among nuclear-encoded genes analyzed.

**Figure 3.**
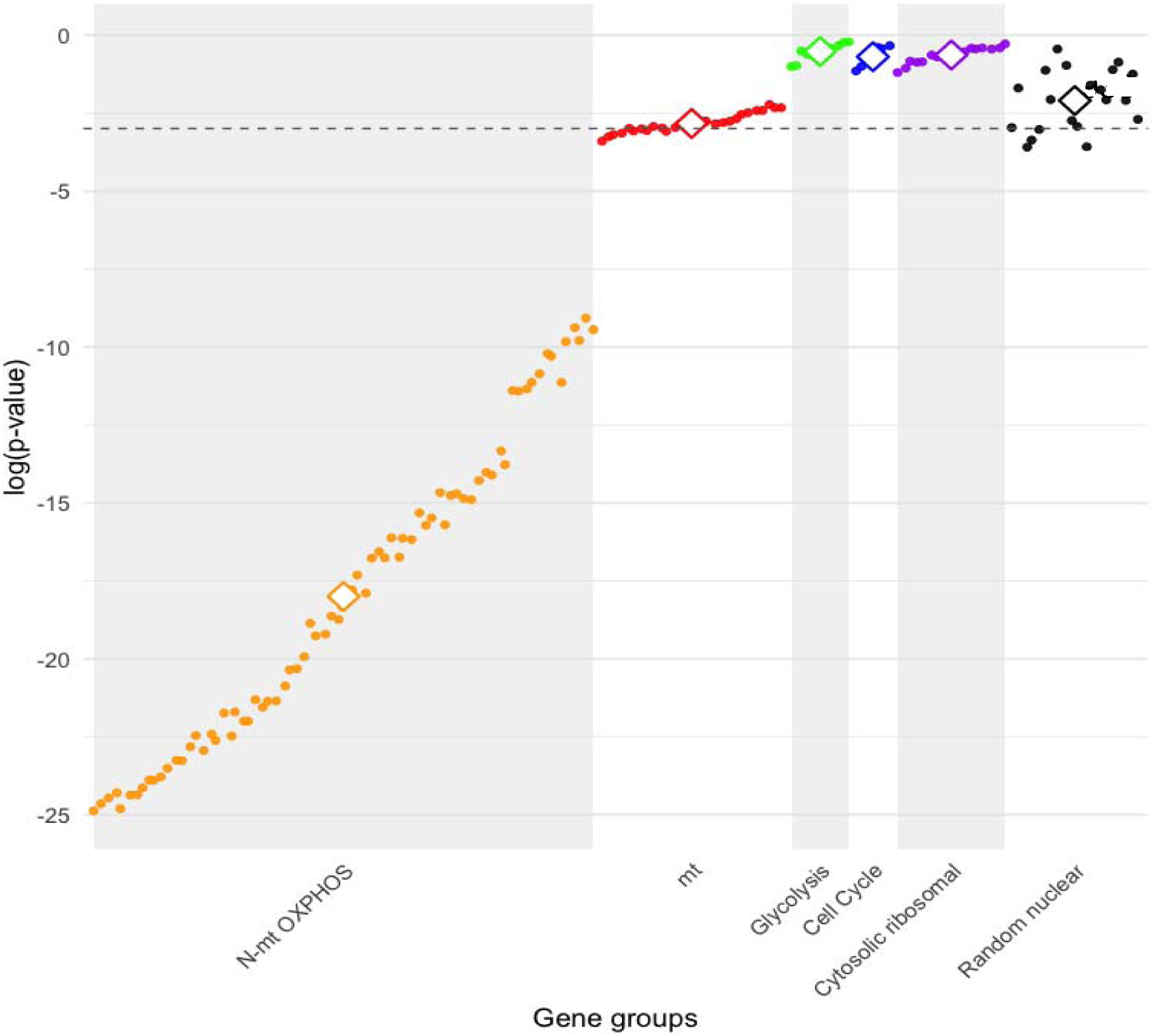
Likelihood ratio test results comparing *d_N_/d_S_* ratios in the 50-kb inversion clade vs. other legumes for all genes in all groups. The null model constrained the entire phylogeny to a single *d_N_/d_S_*ratio, while the alternative model allowed *d_N_/d_S_* to vary between the 50-kb inversion clade and other legumes. Larger squares with white space inside indicate results from the concatenated dataset for each gene group. The horizontal dashed line shows the false discovery rate-corrected 5% significance cutoff, with log *p* values on the y-axis and gene groups ordered on the x-axis. Genes appearing to the left of the vertical dashed line have significantly higher *d_N_/d_S_*in the 50-kb inversion clade relative to other legumes.

We also asked whether *d_N_/d_S_* ratios in N-mt OXPHOS genes were elevated compared to other nuclear-encoded gene groups within the 50-kb inversion clade. *d_N_/d_S_* ratios in the 50-kb inversion clade were generally higher in N-mt OXPHOS genes compared with other nuclear-encoded genes, although this was only statistically significant when comparing N-mt OXPHOS genes to N-cr genes and random nuclear proteins (*P* < 0.001, Wilcoxon rank sum test; *P* > 0.032 for other comparisons).

Elevated *d_N_/d_S_* ratios can be caused by intensified positive selection to fix beneficial mutations or relaxed purifying selection that allows slightly deleterious mutations to fix. Using the program RELAX (Wertheim et al. 2015), we explicitly tested these alternative causes of elevated *d_N_/d_S_* ratios in the 50-kb inversion clade for the different gene groups. Among the mt genes, 82% exhibited *k* values significantly lower than 1.0, and the concatenated set of mt genes showed a similar pattern (*k* = 0.83, *P* = 0.001), consistent with relaxed purifying selection in the 50kb-clade compared to other legumes (Tables S2, S3; Figure 4). Similarly, for N-mt OXPHOS genes, *k* values ranged from 0.502 to 1.831, but 88% of genes and the concatenated gene set displayed *k* values significantly lower than 1.0 (Tables S2, S3; Figure 4), reflecting relatively relaxed purifying selection in the 50-kb inversion clade. However, for all the other nuclear-encoded gene sets, *k* values were generally not significantly different than 1.0, including the concatenated gene set for the N-gly, N-cc, N-cr, and N-rand gene groups (Tables S2, S3; Figure 4), suggesting no differences in selective pressures between the 50-kb inversion clade and other legumes for these datasets.

**Figure 4.**
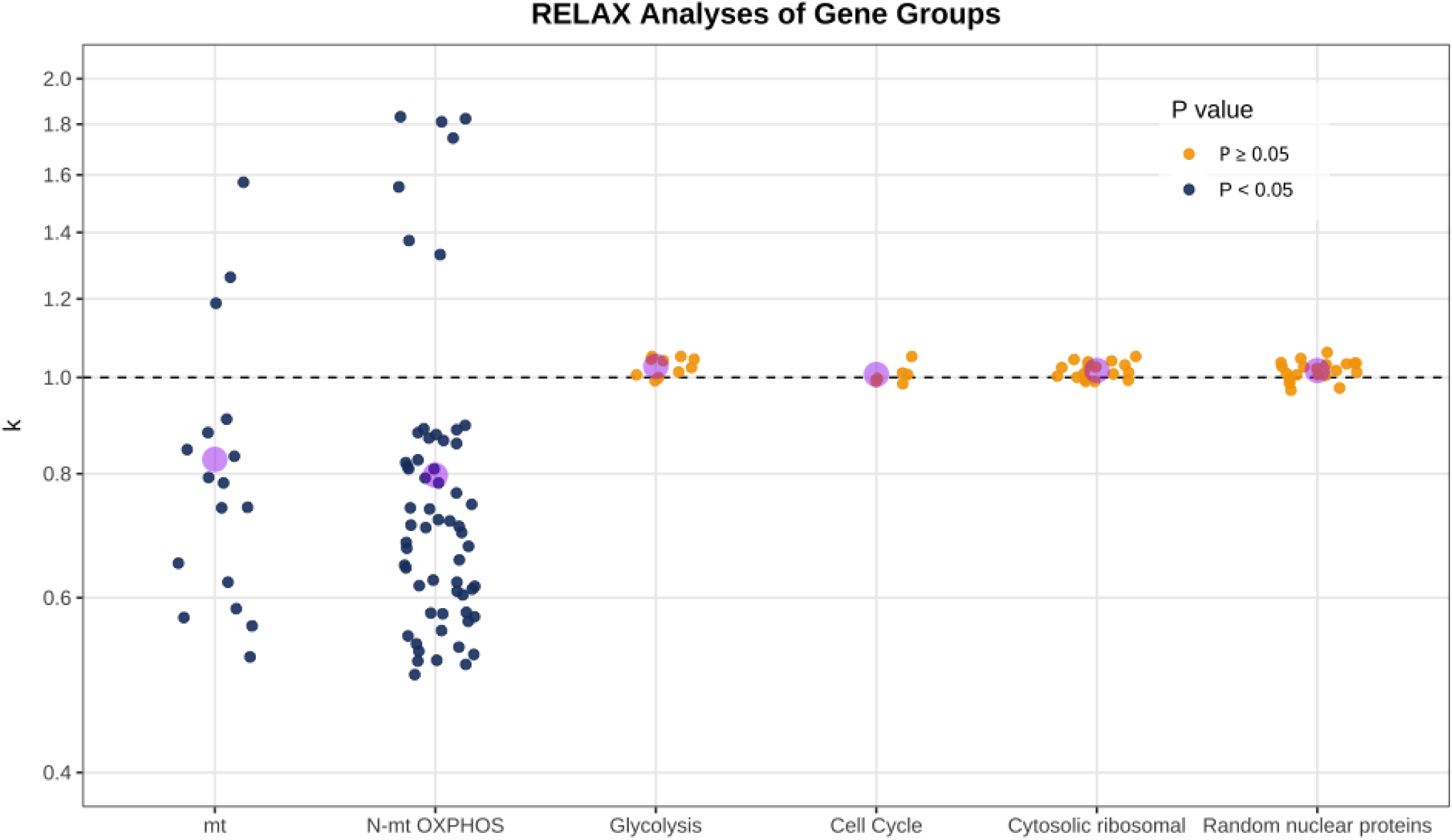
Selection intensity parameter (*k*) estimated for individual genes in six functional categories: mitochondrial (mt), nuclear-encoded mitochondrial oxidative phosphorylation (N-mt OXPHOS), Glycolysis, Cell Cycle, Cytosolic ribosomal, and random nuclear protein genes. Each point represents a single gene, and each gene is colored based on whether the *P*-value was above or below 0.05. Purple circles indicate *k* values from RELAX analyses of concatenated gene datasets. *k* values < 1 suggest relaxed selection in the 50-kb inversion clade compared to other legumes, while values > 1 suggest intensified positive selection in the 50-kb inversion clade.

### Evolutionary rates of mitochondrial genes covary with nuclear OXPHOS genes, but not other nuclear-encoded genes

Evolutionary rate covariation analyses tended to show significant, positive correlations between evolutionary rates in mt and N-mt OXPHOS genes, but no significant correlation between mt rates and nuclear rates for the other gene groups, including N-gly, N-cc, and N-cr genes (Table S4, Figure 5A, Figure S1). Specifically, ERC analyses showed significant positive correlations between branch lengths of mt and N-mt OXPHOS genes in the 50-kb inversion clade (*r_s_* = 0.67, *P* < 0.001 based on bootstrap analyses, Figure 5A, Figure S1). Correlations were not significantly different from zero between mt and nuclear genes in the other gene sets within the 50-kb inversion clade (*P* > 0.5 for all, Figure 5A, Figure S1). When only using taxa outside of the 50-kb inversion clade, no correlations between mt genes and nuclear genes were significant for any gene class (Figure S1). When including all taxa (within and outside of the 50-kb inversion clade), results mirrored those found when using just the 50-kb inversion clade (Figure S1). Additionally, mitochondrial genes and plastid genes showed strong, positive ERC within the 50-kb inversion clade (*r_s_* = 0.89, *P* < 0.001) and across all taxa (Figure 5A, Figure S1E), but not when confined to taxa outside of the 50-kb inversion clade (*r_s_* = -0.35, *P* = 0.36). Results were qualitatively similar when using terminal branch lengths instead of root-to-tip distances (Figure S2).

**Figure 5.**
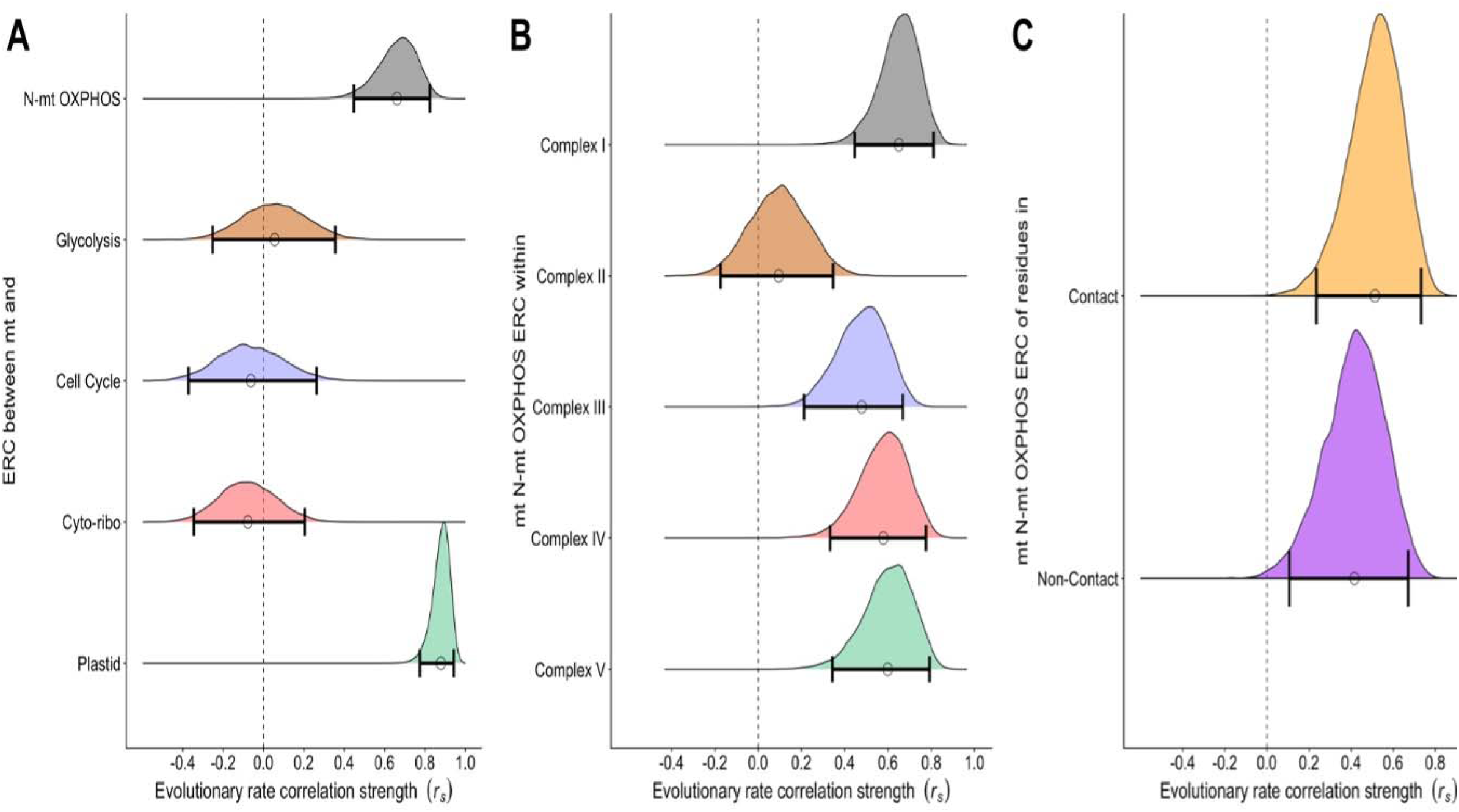
Evolutionary rate covariation (ERC) in the 50-kb inversion clade among mitochondrial (mt) genes and (A) nuclear-encoded genes targeted to the mitochondria (N-mt OXPHOS), non-mitochondrial targeted nuclear-encoded genes (Glycolysis, Cell Cycle, and Cyto-ribo), and plastid genes. The distribution of 10,000 bootstrap replicates of the correlation coefficient, *r_s_* are shown with dots showing the mean and bars showing the 95% confidence intervals. (B) Within-complex ERC of mt and N-mt OXPHOS genes. (C) ERC of mt and N-mt OXPHOS genes that either contain (contact) or lack (non-contact) residues that directly contact mt residues.

When performing ERC between mt-encoded genes and N-mt genes in each OXPHOS complex individually, significant positive correlations between mt and N-mt OXPHOS genes were found across most OXPHOS complexes within the 50-kb inversion clade (Figure 5B, Figure S3), but not in taxa outside the 50-kb inversion clade. Correlations were not statistically different among complexes I, III, IV, and V. However, Complex II, which is composed entirely of nuclear-encoded subunits in Papilionoideae, showed a statistically non-significant correlation within the 50-kb inversion clade (*r_s_* = 0.09, *P* = 0.519, Figure 5B, Figure S3B), and was statistically different than the other complexes based on bootstrap analyses (*P* = 0.001). Similar results were obtained using only terminal branches for ERC analyses (Figure S4).

Narrowing the N-mt OXPHOS dataset to include only complexes with mt-encoded proteins (i.e., excluding Complex II) and partitioning remaining N-mt genes into “contact” and “non-contact” categories (those with or without residues predicted to contact mt-encoded residues) revealed slightly stronger ERC values for genes encoding contact proteins compared to non-contact proteins (*r_s_* = 0.51 vs *r_s_* = 0.41). However, the difference between the two categories was not statistically significant via bootstrap analyses (*P* = 0.507; Table S4, Figure 5C).

## Discussion

### Elevated substitution rates in mitochondrial and nuclear-encoded OXPHOS genes in the 50-kb inversion clade – mitonuclear coevolution or relaxed selection?

In bilaterian mammals, mt genes have been observed to evolve at rates approximately 2 to >20 times faster than nuclear-encoded genes, with the rate varying by species (Allio et al. 2017; Weaver et al. 2022). In most angiosperms, the opposite pattern is usually observed, where the nuclear rate is 15-20 times faster than the mitochondrial genome (Cole et al. 2018). However, in several independent angiosperm lineages, including *Ajuga*, *Acorus*, *Viscum*, *Eleocharis*, *Sarcophyrnium*, *Silene*, *Plantago*, and *Pelargonium*, accelerated mitogenome substitution rates have resulted in patterns resembling those observed in bilaterian mammals (Parkinson et al. 2005; Bakker et al. 2006; Mower et al. 2007; Sloan et al. 2012; Zhu et al. 2014; Skippington et al. 2015; Sloan et al. 2015; Lee et al. 2023). Here, we describe a similar pattern for the 50-kb inversion clade of Papilionoideae legumes, which have significantly accelerated mt *d_S_* values compared to other legumes (Figure 2B). Across gene sets, nuclear-encoded genes generally retain higher *d_S_* than mitochondrial genes, although the difference varies among individual species, from mt *d_S_* being 5.07-fold slower to 1.36-fold higher than nuclear genes based on concatenated alignments and the N-rand dataset (Table S5). More importantly, mt *d_S_* varies dramatically in the 50-kb inversion clade compared to the other legumes we examined, with mt *d_S_*varying by 7.0-fold among species in the 50-kb inversion clade, but only 2.3-fold among other legumes. Although not as drastic, nuclear-encoded genes have also experienced a general rate acceleration in *d_S_* in the 50-kb inversion clade (Figure 2B). Overall, this suggests an increase in mutation rates in the 50-kb inversion clade compared to other legumes, with the mtDNA being especially affected. This is mirrored by significant *d_S_* accelerations of the 50-kb inversion clade in a previous plastid legume study that investigated taxa within and outside of the 50-kb inversion clade (Schwarz et al. 2017).

Both mt and N-mt OXPHOS genes within the 50-kb inversion clade exhibited significantly elevated *d_N_* compared to other legumes, which was not the case for any other gene groups (Figure 2; *P* > 0.239 for other gene groups). The increase in *d_N_*suggests functional consequences and could be indicative of mitonuclear coevolution in the 50-kb inversion clade instead of demographic processes affecting evolutionary rates overall. Changes to mtDNA repair, recombination, or replication systems in the 50-kb inversion clade, as observed in other plant mitochondrial genomes (Mower et al. 2007), could have led to increased mtDNA mutation rates in the 50-kb inversion clade, driving selection for compensatory changes in N-mt genes according to the “nuclear compensation hypothesis”. This hypothesis is a particularly prominent version of mitonuclear coevolution where nuclear-encoded genes are selected for compensatory changes to counteract slightly deleterious mutations accumulating in the mitochondrial genome (Levin et al. 2014; Havird et al. 2017; Sloan et al. 2017; Iannello et al. 2019; Hill 2020).

Under nuclear compensation, we would expect increased *d_N_/d_S_* ratios in the 50-kb inversion clade for N-mt genes, reflecting positive selection for fixing compensatory beneficial mutations. We might also expect increased *d_N_/d_S_* ratios in mt genes, but in this case reflecting relaxed selection where slightly deleterious mutations go to fixation. In the 50-kb inversion clade, we observed significantly elevated *d_N_/d_S_* ratios, but only in N-mt genes (Figure 2C), as predicted under the nuclear compensation hypothesis. Moreover, additional analyses showed all N-mt OXPHOS genes had elevated *d_N_/d_S_* ratios in the 50-kb inversion clade, along with some mt genes, but no other gene groups (Figure 3), providing additional support for the nuclear compensation hypothesis.

However, mt-specific processes could also result in elevated *d_N_* and *d_N_/d_S_* ratios in mt and N-mt genes, including positive or relaxed selection on overall mt function in the 50-kb inversion clade. To be explicit: nuclear compensation predicts elevated *d_N_/d_S_*ratios caused by positive selection in N-mt genes, while changes in selection on overall mt function predicts elevated *d_N_/d_S_* ratios in mt and N-mt genes caused by either positive or relaxed selection. Using tests to explicitly disentangle whether elevated *d_N_/d_S_* ratios in the 50-kb inversion clade were caused by relaxed or positive selection, we found clear evidence of generally relaxed purifying selection acting within both N-mt and mt genes in the 50-kb inversion clade (Figure 4; Table S2), contrary to expectations under nuclear compensation or mitonuclear coevolution in general. While some individual genes did show signs of positive, intensified selection leading to high *d_N_/d_S_* ratios, they were generally spread across both genomes, contrary to nuclear compensation, which would predict positive selection focused on N-mt genes.

While this pattern may reflect unique, unexpected mitonuclear coevolution dynamics associated with individual genes within this lineage, we favor the interpretation that accelerated rates of evolution in mt and N-mt genes in the 50-kb inversion clade are due to relaxed selection on mt processes in the 50-kb inversion clade and not indicative of mitonuclear coevolution. Functional analyses of mitochondrial processes in species in the 50-kb inversion clade compared to other species could more directly reveal altered mt function, as has been documented in some plants with accelerated mt evolution (Maclean et al. 2018; Weaver et al. 2020; Cai et al. 2025). In contrast, studies on *Silene* species demonstrated that rapid mitochondrial genome evolution most likely led to compensatory evolution in nuclear-encoded mitochondrial genes, with no evidence for relaxed selection and some evidence for positive selection in N-mt genes using similar tests (Havird et al. 2017). Elevated substitution rates in *Silene* N-mt genes also often occur at residues that interact with mitochondrial-encoded residues, supporting their role in maintaining OXPHOS complex stability under increased mutational pressure (Havird et al. 2015). Similarly, research on *Pelargonium* has shown that both mitochondrial and plastid genes exhibit accelerated substitution rates, with mitochondrial genes displaying elevated synonymous substitution rates and plastid genes showing increased nonsynonymous rates in specific protein-coding regions (Weng et al. 2012). Our findings highlight how lineage-specific genomic dynamics can drive elevated rates of nucleotide substitutions and contribute to the evolutionary complexity of organellar genomes. While coordinated evolution between mitochondrial and nuclear genomes is undoubtedly important, accelerated rates in mt and N-mt genes should be explored carefully before they are assumed to be a signature of mitonuclear coevolution.

### Despite relaxed selection, evolutionary rate correlations suggest a role for mitonuclear coevolution

Evolutionary rate covariation (ERC) analyses revealed a general pattern where mt rates were positively correlated with N-mt rates in the 50-kb inversion clade but not rates of other nuclear-encoded genes (Figure 5A), which would be predicted if demographic processes were driving variation in evolutionary rates. This pattern has been found across animals (Yan et al. 2019; Piccinini et al. 2021; Weaver et al. 2022; Smith et al. 2024; Tao et al. 2024; Wallnoefer et al. 2025) and has generally been interpreted as positive evidence for mitonuclear coevolution. However, shared evolutionary constraints on similar functions encoded by different genes, including mitochondrial genes in different genomes, could also produce strong ERC. Here, we observed stark differences in mitonuclear ERC across OXPHOS complexes, with those from Complex II showing no correlation with rates of mt genes (Figure 5B). Complex II is entirely nuclear-encoded in many taxa, including legumes, and serves as a negative control for mitonuclear interactions and processes driven by mitonuclear coevolution. For example, experimental studies in *Tigriopus* copepods and *Xiphophorus* fishes show that mitonuclear mismatches in hybrids reduce OXPHOS function, but Complex II remains unaffected (Burton 1990; Ellison and Burton 2006; Barreto et al. 2015; Moran et al. 2024; Robles et al. 2025). Among the other electron transport chain complexes, Complex I has the highest number of mitochondrial-encoded genes and showed the strongest correlation with mt rates (Figure 5B). Moreover, within the electron transport system complexes, nuclear-encoded proteins that closely interact with mitochondrial-encoded subunits exhibited the strongest correlations compared to proteins that do not directly contact mt-encoded residues (Figure 5C, although not reaching statistical significance).

Together, although we find evidence for shared relaxation of selection on mt and N-mt genes, the strong evolutionary rate correlations reported here in the 50-kb inversion clade between mt and N-mt OXPHOS genes with direct mt contacts appear to be based at least in part on mitonuclear interactions and not shared mitochondrial function. This phenomenon supports a role for mitonuclear coevolution in OXPHOS gene evolution and has been broadly supported across taxa, from mammals to plants, where mitonuclear interactions play a crucial role in shaping the evolution of oxidative phosphorylation complexes (Gershoni et al. 2010; Levin et al. 2014; Sloan et al. 2017; Hill et al. 2019). Although somewhat contradictory to our findings above, we hypothesize that while relaxed selection may have led to overall higher mt rates in the 50-kb inversion clade, this also led to more variation in mt rates in the 50-kb inversion clade. Similar to mt *d_S_*, terminal branch lengths were uniformly low in legumes outside of the 50-kb inversion clade (varying ∼2-fold), but within the 50-kb inversion clade, mt branch lengths varied widely (varying ∼22-fold, although somewhat driven by outliers). This variation may have driven selection for complementary changes in some species within the 50-kb inversion clade, or in particular genes. This variation may also provide the statistical power needed to detect significant ERC in the 50-kb inversion clade, while uniformly low mt rates limited our ability outside of the 50-kb inversion clade.

Overall, our results suggest a complicated view of OXPHOS evolution compared to previously described taxa. Widespread elevated rates of evolution in the 50-kb inversion clade are likely not reflective of mitonuclear coevolution, even though *d_N_/d_S_* ratios were only statistically elevated in N-mt genes in some analyses, as predicted under nuclear compensation. Rather, relaxed selection on mitochondrial processes explains elevated rates in the 50-kb inversion clade compared to other legumes for most OXPHOS genes. However, ERC analyses point to physical interactions between mt and nuclear-encoded OXPHOS genes as important drivers of rate covariation within the 50-kb inversion clade. Although complex, this suggests both shared functional constraints and coevolution explain evolutionary rate variation in this system. Other studies on mitonuclear interactions should consider both of these processes when examining evolutionary rates in mt- and nuclear-encoded genes.

### Similar, but different, evolutionary pressures shape mitochondrial- and plastid-nuclear interactions in the 50-kb inversion clade

A strong correlation was observed between mitochondrial and plastid branch lengths in the 50-kb inversion clade (Figure 5A, Figure S1E), similar in strength to the ERC between mt and N-mt genes. Despite some recent evidence suggesting physical interactions between mitochondria and plastids may be possible in plant and diatom cells (Oikawa et al. 2021; He et al. 2023; Giustini et al. 2025), it is unlikely that plastid and mt genes are under selection to coevolve with each other. Similarly, plastid and mitochondrial genes participate in different cellular processes (largely oxidative phosphorylation vs. photosynthesis), so it is unlikely that shared evolutionary constraints on a similar function are responsible for ERC between mt and plastid genes. Demographic changes are not likely responsible either, as mt rates were not correlated with nuclear rates in most nuclear control gene groups (Figure 5). Rather, we suggest shared mechanisms in DNA homeostasis may underly the shared evolutionary rates across mt and plastid genes in the 50-kb inversion clade. Many nuclear-encoded proteins responsible for DNA repair, replication, and recombination are dual-targeted to both plastids and mitochondria in land plants (Morley et al. 2019). Changes in these dual-targeted nuclear genes could lead to similar rate changes in both mt and plastid genomes. Furthermore, while *d_S_* were not elevated in plastid-encoded ribosomal genes in the 50-kb inversion clade, *d_N_* was accelerated (Tressel et al. 2025) and the key molecular synapomorphy of a 50-kb inversion in the plastid genome in this clade also suggests altered genome maintenance in the plastid genome.

Our previous work investigating coevolution between plastid and nuclear genomes revealed similar patterns as those described here for mitonuclear coevolution (Tressel et al. 2025). Most notably, rate correlations were strong between plastid-encoded genes and their interacting nuclear-encoded genes, but only for the 50-kb inversion clade, not other legumes. *d_N_* and *d_N_/d_S_* ratios were also elevated for plastid-encoded and their interacting nuclear-encoded genes in the 50-kb inversion clade compared to other legumes, but not for other nuclear-encoded gene groups. However, unlike mt genes, no plastid-related genes showed elevated *d_S_* in the 50-kb inversion clade. Taken together, we suggest overall similar evolutionary rate changes in both mitochondrial and plastid genomes of the 50-kb inversion clade could be caused by using similar DNA repair, replication, and recombination machinery, leading to strong signals of cytonuclear coevolution for both mitochondrial and plastid genes. Similarly, in *Silene,* Geraniaceae, Plantaginaceae, and *Eleocharis* mt rate accelerations are mirrored by plastid rate accelerations and some evidence for cytonuclear coevolution in both genomes (Parkinson et al. 2005; Bakker et al. 2006; Mower et al. 2007; Havird et al. 2015, 2017; Rockenbach et al. 2016; Lee et al. 2023), although only a subset of plastid genes may be affected. While the specifics may be different in different taxa and in plastid vs. mitochondrial genomes, cytonuclear coevolution is likely an important feature of organelle genomes in general, with plants offering a powerful system to explore and disentangle the processes that underly cytonuclear coevolution.

## Conclusions

Coordination between nuclear and mitochondrial genomes is essential for maintaining mitochondrial function and supporting key processes including energy production and stress response. Here, we uncover signatures of mitonuclear coevolution within the legume 50-kb inversion clade, with strong evolutionary rate correlations between mitochondrial-encoded and nuclear-encoded mitochondrial-targeted genes. Unlike in previously examined taxa, we find that the wide variation in mt evolutionary rates in this clade is likely driven by overall relaxed selection on OXPHOS genes compared to other legumes. Broader questions remain about how coevolutionary dynamics between mt and nuclear genomes compare to those between plastid and nuclear genomes, both within legumes and across plants in general. While relatively low mitochondrial mutation rates in angiosperms may obscure the accumulation of mitonuclear incompatibilities and signals of mitonuclear coevolution, comparative studies across diverse taxa are needed to elucidate general principles cytonuclear coevolution. By expanding our understanding of the mechanisms driving cytonuclear interactions, particularly in agriculturally important clades like Papilionoideae, this research provides a foundation for exploring how genomic coordination underpins plant adaptation and evolutionary resilience.

## Materials and Methods

### Taxon sampling, DNA/RNA isolation, sequencing, and transcriptome assembly

Our dataset included 50 species: 48 Papilionoideae (including 41 from the 50-kb inversion clade), one Cercidoideae, and one Caesalpinioideae (Table S6). Together, the papilionoid species represented 15 of the 22 early diverging major clades identified in phylogenetic studies of the subfamily (Cardoso et al. 2012, 2013; Choi et al. 2022; Cai et al. 2025). Previously sequenced genomic DNA (Lee et al. 2021; Choi et al. 2022), was used to draft 48 assembled mitogenomes, and paired with transcriptomes generated for each taxon. To expand sampling, two additional transcriptomes and one published mitogenome were incorporated, while raw genomic reads from NCBI were used to assemble an additional draft mitogenome (Table S7). Transcriptomes, originally generated in Tressel et al. (2025), were sourced from plant materials either collected in the field or cultivated in the greenhouse at The University of Texas at Austin (UT-Austin). Voucher specimens were archived at the Billie L. Turner Plant Resources Center (TEX/LL) at UT-Austin, the Rio de Janeiro Botanical Garden Herbarium (RB), and the Herbarium at the Universidade Estadual de Feira de Santana (HUEFS) (Table S6). RNA isolation and sequencing for data not downloaded from NCBI was described previously in Tressel et al. (2025). Transcriptome assembly for all 50 species including quality control, adapter trimming, rRNA read removal, *de novo* assembly, assembly quality and completeness assessments are also described in Tressel et al. (2025; Table S1).

### Mitochondrial-encoded and nuclear-encoded gene curation

Fifty draft mitogenomes were utilized, including one downloaded mitogenome and another assembled from downloaded raw reads both from NCBI. Genome assembly, read mapping, and annotations followed procedures previously described in Choi et al. (2019). Relevant mitochondrial-encoded OXPHOS gene sequences for our analyses were extracted from these annotations. Transcriptome assemblies were searched for nuclear-encoded OXPHOS genes using tBLASTn (version 2.12.0+) with *Arabidopsis thaliana* protein sequences from TAIR (Berardini et al. 2015) as queries. The total number of nuclear- and mitochondrial-encoded OXPHOS genes analyzed in each OXPHOS complex was as follows: Complex I: 29, 9; Complex II: 7,0; Complex III: 9, 1; Complex IV: 6, 2; Complex V: 9, 5. We also extracted nuclear-encoded control gene sets using a similar approach, totaling 53 nuclear-encoded genes: 10 from the glycolysis pathway (N-gly), 6 from the cell cycle pathway (N-cc), 17 from cytosolic ribosomes (N-cr), and 20 haphazardly chosen nuclear genes (N-rand, Yates et al. 2022; Table S8).

### Dataset compilation and alignment construction

For nuclear-encoded, mitochondrial-targeted OXPHOS genes (N-mt), N-terminal targeting peptides were predicted using the TargetP webserver (https://services.healthtech.dtu.dk/services/TargetP-2.0/, last accessed on June 15, 2024, Emanuelsson et al. (2007) and removed prior to alignment. Sequences were aligned for each gene using MAFFT v7.490 (Katoh and Standley 2013) and all gene alignments within a gene group were also concatenated in Geneious v. 2023.1.1 (https://www.geneious.com; Kearse et al. 2012)

### Constraint tree construction

Phylogenetic analysis was conducted on the RAxML web server (Stamatakis et al. 2019) with default parameters to generate a maximum likelihood tree. The dataset used consisted of 59 concatenated plastid-protein coding genes without partitioning (Table S1). To evaluate support for clades, 1000 bootstrap replicates were performed. In addition, 100 independent runs were conducted, and the highest-scoring maximum likelihood (ML) tree was selected. This resulting topology was subsequently used as the constraint tree for further substitution rate estimate and coevolution analyses.

### Estimating d_N_, d_S_, and d_N_/d_S_

Nucleotide sequence alignments of individual genes in the six gene groups were used to estimate *d_N_, d_S_,* and *d_N_/d_S_* in the 50-kb inversion clade vs. other legumes with HyPhy and codeml (model = 2) in PAML 4.9j (Pond et al. 2005; Yang 2007; Table S9) by using methods and scripts described in Tressel (2025). Briefly, a custom bash script with the codon model MG94xHKY85 in HyPhy (Pond et al. 2005) was run assigning branches with thick lines in Figure 1 one rate for the 50-kb inversion clade while all other branches were assigned another rate for other legumes as the alternative model and assumed a shared *d_N_* and *d_S_* for the null model. The *d_N_* and *d_S_* values for the 50-kb inversion clade and other legumes were summed and binned by gene group. To test for rate differences, we used Wilcoxon rank sum tests to compare rates in the 50-kb inversion clade to rates for other legumes for each gene group and N-mt OXPHOS genes to other nuclear-encoded gene groups within the 50-kb inversion clade. Resulting *P* values were adjusted using false discovery rate (FDR).

We also used likelihood ratio tests (LRTs) to compare rates between the 50-kb inversion clade and other legumes across each gene in the different gene groups using codeml in PAML 4.9j (Yang 2007). In this analysis, the null model (model = 0) assumed a shared *d_N_/d_S_*across the entire phylogenetic tree, while the alternative model (model = 2) allowed for different *d_N_/d_S_* in the 50-kb inversion clade versus the other legume lineages. We also applied this LRT approach to the concatenated set of all genes within each of the gene groups.

### Detection of relaxed vs. intensified positive selection

Relaxed purifying selection or intensified positive selection was tested for in the 50-kb inversion clade compared to the other taxa using the RELAX program (Wertheim et al. 2015) implemented in HyPhy (Pond et al. 2005) for individual genes and the concatenated set within each gene group. The 50-kb inversion clade (thick branches in Figure 1) was designated as the test branch, while the remaining Fabaceae lineages served as the reference branches. RELAX incorporates a selection intensity parameter (*k*) to describe how categories of sites based on *d_N_/d_S_* differs between the test and reference branches (either tending to converge or diverge from *d_N_/d_S_* = 1). In the null model, *k* is fixed at 1.0, representing no difference in selection intensity, whereas in the alternative model, *k* is allowed to vary. A likelihood ratio test is used to determine if *k* significantly deviated from 1.0. A rejection of the null model, coupled with higher *d_N_/d_S_* values in the test branches and *k* values less than 1.0, would indicate that purifying selection was relaxed in the 50-kb inversion clade compared to the other Fabaceae lineages, while *k* > 1 would suggest intensified positive selection in the 50-kb inversion clade.

### Contact residue identification

The CyMIRA database (Forsythe et al. 2019), a comprehensive resource for characterizing cytonuclear molecular interactions in plants, was used to identify contact residues in N-mt OXPHOS genes that physically contact residues encoded by mt genes. CyMIRA provides residue-level data on direct cytonuclear contact sites derived from curated structural and interaction datasets. Sites were classified as “contact” in N-mt OXPHOS genes if they contacted at least one mt residue. N-mt OXPHOS genes lacking any contact mt residues were considered “non-contact”, while those with a contact were considered “contact” genes.

### Evolutionary rate covariation (ERC) analyses

Evolutionary rates for each set of concatenated nucleotide sequence alignments (Table S4) were assessed by estimating branch lengths on a ML phylogeny using the RAxML web server (Stamatakis et al. 2019). The topology was constrained by the plastid protein-coding species tree, fixing it according to the known relationships within Fabaceae. Branch lengths were estimated using the best-fitting nucleotide substitution model, with 1000 bootstraps followed by ML search and optimization. Root-to-tip branch lengths were extracted using the distRoot function within the “adephylo” and “ape” packages in R (Paradis et al. 2019). Terminal branch lengths were also extracted using a custom R function that isolates the edge length associated with each terminal tip from the phylogenetic tree’s edge matrix. Phylogenies were additionally generated for each OXPHOS complex individually, based on concatenated sets of mitochondrial and nuclear-encoded mitochondrial genes.

Evolutionary rate covariation (ERC) analysis was used to test whether rates of evolution in the mitogenome were more highly correlated with rates of evolution in interacting nuclear-encoded genes or ones that lack mitochondrial interactions using mt, N-mt OXPHOS (all together and contact vs. non-contact genes considered separately), N-gly, N-cc, N-cr, and plastid (17 plastid-encoded ribosomal genes used in Tressel et al. 2025) (Table S9) gene sets. To avoid spurious correlations among gene sets that are not coevolving, each target gene set was normalized to the background evolutionary rate by dividing the branch lengths of each target gene set with a random set of 20 nuclear proteins (split into two sets of 10 genes each, one used for the mt genes and one used for the nuclear genes) selected using EnsemblPlants (Yates et al. 2022; Table S8). To estimate evolutionary rate correlations, Spearman rank correlation tests on the normalized branch lengths of each gene set were conducted using *corr.test* in R (version 4.1.2, R Core Team 2021). For evolutionary rate correlations within OXPHOS complexes, mt and N-mt OXPHOS genes within that complex were compared. However, CII has no mt-encoded genes, so the overall mt gene dataset was used for that evolutionary rate correlation analyses. Spearman rank correlation estimates were bootstrapped (10,000 iterations) to test for statistically different correlations (*r_s_*) within gene sets using the boot function in R. The difference between the *r_s_* distribution of each gene set was calculated. From the resulting *r_s_* distribution, 95% confidence intervals were determined using the *boot.ci* function in R, with the type set to “bca”. When 95% confidence intervals did not overlap zero, we considered this a significant ERC, and if 95% confidence intervals did not overlap the mean *r_s_* of a different gene set, we considered the gene sets to have significantly different *r_s_* distributions.

## Acknowledgements

We thank the TEX-LL, HUEFS, and RB herbaria for voucher deposition, the Desert Legume Program at the University of Arizona for seeds. We also thank George Yatskievych (TEX/LL) for arranging a formal Material Transfer Agreement (Decree number 8772) under the SisGen Cadastro RDC6BE9, which facilitated research activities between our institutions. Finally, we thank Luciano P. de Queiroz (HUEFS) and Haroldo C. de Lima (RB) for providing access to living collections and for arranging field work in Brazil.

## Funding sources list

This work was supported by grants from the National Science Foundation (DEB-1853010 and DEB-1853024) to MW, RJ, and TR, the Texas Ecological Laboratory Program (EcoLab) to RJ, TR, and I-SC, the Sidney F. and Doris Blake Professorship in Systematic Botany to RJ, the University of Texas at Austin Graduate School Continuing Fellowship to LGT, NIH R35GM142836 and EDGE–NSF IOS-2421661 to JCH, the CNPq (Research Productivity Fellowship no. 308244/2018-4; Universal no. 422325/2018-0), FAPESB (Universal no. APP0037/2016), and UFBA PROQUAD program to DC.

## Data availability

All raw RNA sequencing data have been deposited in the NCBI GenBank repository under Bioproject PRJNA1120003. All mitochondrial CDS sequence accessions are shown in Table S7 and have been deposited in NCBI GenBank. All scripts used in this study are available on GitHub: https://github.com/Ltressel/Cytonuclear_Coevolution_Relaxed_Selection.

## References

Allio R, Donega S, Galtier N, Nabholz B. 2017. Large variation in the ratio of mitochondrial to nuclear mutation rate across animals: implications for genetic diversity and the use of mitochondrial DNA as a molecular marker. Mol. Biol. Evol. 34:2762–2772.

Alverson AJ, Zhuo S, Rice DW, Sloan DB, Palmer JD. 2011. The mitochondrial genome of the legume *Vigna radiata* and the analysis of recombination across short mitochondrial repeats. PLoS One. 6:e16404.

Asar Y, Sauquet H, Ho SYW. 2024. Evolutionary rates of nuclear and organellar genomes are linked in land plants. bioRxiv, unpublished data, 10.1101/2024.08.05.606707.

Bakker FT, Breman F, Merckx V. 2006. DNA sequence evolution in fast-evolving mitochondrial DNA nad1 exons in Geraniaceae and Plantaginaceae. Taxon. 55:887–896.

Barreto FS, Pereira RJ, Burton RS. 2015. Hybrid dysfunction and physiological compensation in gene expression. Mol Biol Evol. 32:613–622.

Berardini TZ, Reiser L, Li D, Mezheritsky Y, Muller R, Strait E, Huala E. 2015. The Arabidopsis Information Resource: making and mining the “gold standard” annotated reference plant genome. Genesis. 53:474–485.

Bi C, Wang X, Xu Y, Wei S, Shi Y, Dai X, Yin T, Ye N. 2016. The complete mitochondrial genome of *Medicago truncatula*. Mitochondrial DNA B. 1:122–123.

Burton RS. 1990. Hybrid breakdown in developmental time in the copepod *Tigriopus californicus*. Evolution. 44:1814–1822.

Burton RS, Pereira RJ, Barreto FS. 2013. Cytonuclear genomic interactions and hybrid breakdown. Annu Rev Ecol Evol Syst. 44:281–302.

Cai L, Cardoso D, Tressel LG, Lee C, Shrestha B, Choi I-S, de Lima HC, de Queiroz LP, Ruhlman TA, Jansen RK, Wojciechowski MF. 2025. Well-resolved phylogeny supports repeated evolution of keel flowers as a synergistic contributor to papilionoid legume diversification. New Phytol. doi:10.1111/nph.70080.

Cai L, Jansen RK, Havird JC. 2025. Altered mitochondrial respiration is associated with loss of nuclear-encoded OXPHOS genes in parasitic broomrapes. Ecol Evol. 15:e71737. doi:10.1002/ece3.71737.

Cardoso D, de Queiroz LP, Pennington RT, de Lima HC, Fonty É, Wojciechowski MF, Lavin M. 2012. Revisiting the phylogeny of papilionoid legumes: new insights from comprehensively sampled early-branching lineages. Am J Bot. 99:1991–2013.

Cardoso D, Pennington RT, de Queiroz LP, Boatwright JS, van Wyk B-E, Wojciechowski MF, Lavin M. 2013. Reconstructing the deep-branching relationships of the papilionoid legumes. S Afr J Bot. 89:58–75.

Chang S, Wang Y, Lu J, Gai J, Li J, Chu P, Guan R, Zhao T. 2013. The mitochondrial genome of soybean reveals complex genome structures and gene evolution at intercellular and phylogenetic levels. PLoS One. 8:e56502.

Choi I-S, Cardoso D, de Queiroz LP, de Lima HC, Lee C, Ruhlman TA, Jansen RK, Wojciechowski MF. 2022. Highly resolved papilionoid legume phylogeny based on plastid phylogenomics. Front Plant Sci. 13:823190.

Choi I-S, Schwarz EN, Ruhlman TA, Khiyami MA, Sabir JSM, Hajarah NH, Sabir MJ, Rabah SO, Jansen RK. 2019. Fluctuations in Fabaceae mitochondrial genome size and content are both ancient and recent. BMC Plant Biol. 19:448.

Choi I-S, Wojciechowski MF, Ruhlman TA, Jansen RK. 2021. In and out: evolution of viral sequences in the mitochondrial genomes of legumes (Fabaceae). Mol Phylogenet Evol. 163:107236.

Clark NL, Alani E, Aquadro CF. 2012. Evolutionary rate covariation reveals shared functionality and coexpression of genes. Genome Res. 22:714–720.

Cole LW, Guo W, Mower JP, Palmer JD. 2018. High and variable rates of repeat-mediated mitochondrial genome rearrangement in a genus of plants. Mol Biol Evol. 35:2773–2785.

de Juan D, Pazos F, Valencia A. 2013. Emerging methods in protein coevolution. Nat Rev Genet. 14:249–261.

Doyle JJ, Doyle JL, Ballenger JA, Palmer JD. 1996. The distribution and phylogenetic significance of a 50-kb chloroplast DNA inversion in the flowering plant family Leguminosae. Mol Phylogenet Evol. 5:429–438.

Dutta A, Trivedi A, Nath CP, Gupta DS, Hazra KK. 2022. A comprehensive review on grain legumes as climate-smart crops: challenges and prospects. Environ Challenges. 7:100479.

Ellison CK, Burton RS. 2006. Disruption of mitochondrial function in interpopulation hybrids of *Tigriopus californicus*. Evolution. 60:1382–1391.

Emanuelsson O, Brunak S, von Heijne G, Nielsen H. 2007. Locating proteins in the cell using TargetP, SignalP and related tools. Nat Protoc. 2:953–971.

Forsythe ES, Gatts TC, Lane LE, deRoux C, Berggren MJ, Rehmann EA, Zak EN, Bartel T, L’Argent LA, Sloan DB. 2025. ERCnet: phylogenomic prediction of interaction networks in the presence of gene duplication. Mol Biol Evol. 42:1–16.

Forsythe ES, Sharbrough J, Havird JC, Warren JM, Sloan DB. 2019. CyMIRA: the Cytonuclear Molecular Interactions Reference for Arabidopsis. Genome Biol Evol. 11:2194–2202.

Forsythe ES, Williams AM, Sloan DB. 2021. Genome-wide signatures of plastid-nuclear coevolution point to repeated perturbations of plastid proteostasis systems across angiosperms. Plant Cell. 33:980–997.

Giustini C, Dal Bo D, Storti M, Van Vlierberghe M, Baurain D, Cardol P, Zhang Y, Fernie AR, Fitzpatrick D, Aro E-M, et al. 2025. A mitochondrially related plastidial transporter regulates photosynthesis in the diatom *Phaeodactylum tricornutum*. Physiol Plant. 177:e70640. doi:10.1111/ppl.70640.

Gershoni M, Fuchs A, Shani N, Fridman Y, Corral-Debrinski M, Aharoni A, Frishman D, Mishmar D. 2010. Coevolution predicts direct interactions between mtDNA-encoded and nDNA-encoded subunits of oxidative phosphorylation complex I. J Mol Biol. 404:158–171.

Gray MW, Burger G, Lang BF. 1999. Mitochondrial evolution. Science. 283:1476–1481.

Guo X, Wang H, Lin D, Wang Y, Jin X. 2024. Cytonuclear evolution in fully heterotrophic plants: lifestyles and gene function determine scenarios. BMC Plant Biology. 24:989.

Havird JC, Trapp P, Miller C, Bazos I, Sloan DB. 2017. Causes and consequences of rapidly evolving mtDNA in a plant lineage. Genome Biol Evol. 9:323–336.

Havird JC, Whitehill NS, Snow CD, Sloan DB. 2015. Conservative and compensatory evolution in oxidative phosphorylation complexes of angiosperms with highly divergent rates of mitochondrial genome evolution. Evolution. 69:2069–3081.

He C, Berkowitz O, Hu S, Zhao Y, Qian K, Shou H, Whelan J, Wang Y. 2023. Co-regulation of mitochondrial and chloroplast function: molecular components and mechanisms. Plant Commun. 4:100496.

Hill GE. 2015. Mitonuclear ecology. Mol Biol Evol. 32:1917–1927.

Hill GE. 2020. Mitonuclear compensatory coevolution. Trends Genet. 36:403–414.

Hill GE, Havird JC, Sloan DB, Burton RS, Greening C, Dowling DK. 2019. Assessing the fitness consequences of mitonuclear interactions in natural populations. Biol Rev. 94:1089–1104.

Iannello M, Puccio G, Piccinini G, Passamonti M, Ghiselli F. 2019. The dynamics of mito-nuclear coevolution: a perspective from bivalve species with two different mechanisms of mitochondrial inheritance. J Zool Syst Evol Res. 57:534–547.

Katoh K, Standley DM. 2013. MAFFT multiple sequence alignment software version 7: improvements in performance and usability. Mol Biol Evol. 30:772–780.

Kazakoff SH, Imelfort M, Edwards D, Koehorst J, Biswas B, Batley J, Scott PT, Gresshoff PM. 2012. Capturing the biofuel wellhead and powerhouse: the chloroplast and mitochondrial genomes of the leguminous feedstock *tree Pongamia pinnata*. PLoS One. 7:e51687.

Kearse, M. et al. 2012. Geneious Basic: an integrated and extendable desktop software platform for the organization and analysis of sequence data. Bioinformatics 28. 1647–1649.

Kovar L, Nageswara-Rao M, Ortega-Rodriguez S, Dugas DV, Straub S, Cronn R, Strickler SR, Hughes CE, Hanley KA, Rodriguez DN. 2018. PacBio-based mitochondrial genome assembly of *Leucaena trichandra* (Leguminosae) and an intrageneric assessment of mitochondrial RNA editing. Genome Biol Evol. 10:2501–2517.

Lee C, Cardoso D, de Lima HC, de Queiroz LP, Wojciechowski MF, Jansen RK, Ruhlman TA. 2021. The chicken or the egg? Plastome evolution and an independent loss of the inverted repeat in papilionoid legumes. Plant J. 107:861–875.

Lee C, Ruhlman TA, Jansen RK. 2023. Rate accelerations in plastid and mitochondrial genomes of Cyperaceae occur in the same clades. Mol Phylogenet Evol. 182:107760.

Legume Phylogeny Working Group (LPWG). 2017. A new subfamily classification of the Leguminosae based on a taxonomically comprehensive phylogeny. Taxon. 66:44–77.

Levin L, Blumberg A, Barshad G, Mishmar D. 2014. Mito-nuclear coevolution: the positive and negative sides of functional ancient mutations. Front Genet. 5:1–11.

Lian C, Xue Y, Han J, Zhang Y, Zhao H. 2024. Association analysis provides insights into plant mitonuclear interactions. Mol Biol Evol. 41:1532–1543.

Maclean AE, Hertle AP, Ligas J, Bock R, Balk J, Meyer EH. 2018. Absence of complex I is associated with diminished respiratory chain function in European mistletoe. Curr Biol. 28:1614–1619.

Morley SA, Ahmad N, Nielsen BL. 2019. Plant organelle genome replication. Plants. 8:358. doi:10.3390/plants8100358.

Moran BM, Payne CY, Powell DL, Iverson ENK, Donny AE, Banerjee SM, Langdon QK, Gunn TR, Rodriguez-Soto RA, Madero A, et al. 2024. A lethal mitonuclear incompatibility in complex I of natural hybrids. Nature. doi:10.1038/s41586-023-06895-8.

Mower JP, Sloan DB, Alverson AJ. 2012. Plant mitochondrial genome diversity: the genomics revolution. In: Wendel JF, Greilhuber J, Doležel J, Leitch IJ, editors. Plant genome diversity. Vol. 1. Plant genomes. Vienna (Austria): Springer. p. 123–144.

Mower JP, Touzet P, Gummow JS, Delph LF, Palmer JD. 2007. Extensive variation in synonymous substitution rates in mitochondrial genes of seed plants. BMC Evol Biol. 7:135.

Naito K, Kaga A, Tomooka N, Kawase M. 2013. De novo assembly of the complete organelle genome sequences of azuki bean (*Vigna angularis*) using next-generation sequencers. Breed Sci. 63:176.

Negruk V. 2013. Mitochondrial genome sequence of the legume *Vicia faba*. Front Plant Sci. 4:128.

Oikawa K, Imai T, Thagun C, Toyooka K, Yoshizumi T, Ishikawa K, Kodama Y, Numata K. 2021. Mitochondrial movement during its association with chloroplasts in *Arabidopsis thaliana*. Commun Biol. 4:292. doi:10.1038/s42003-021-01833-8.

Paradis E, Schliep K. 2019. APE 5.0: an environment for modern phylogenetics and evolutionary analyses in R. Bioinformatics. 35:526–528.

Parkinson CL, Mower JP, Qiu Y-L, Shirk AJ, Song K, Young ND, de Pamphilis CW, Palmer JD. 2005. Multiple major increases and decreases in mitochondrial substitution rates in the plant family Geraniaceae. BMC Evol Biol. 5:1–12.

Piccinini G, Iannello M, Puccio G, Plazzi F, Havird JC, Ghiselli F. 2021. Mitonuclear coevolution, but not nuclear compensation, drives evolution of OXPHOS complexes in bivalves. Mol Biol Evol. 38:2597–2614.

Pond SL, Frost SDW, Muse SV. 2005. HyPhy: hypothesis testing using phylogenies. Bioinformatics. 21:676–679.

Rand DM, Haney RA, Fry AJ. 2004. Cytonuclear coevolution: the genomics of cooperation. Trends Ecol Evol. 19:645–653.

R Core Team. 2021. R: a language and environment for statistical computing. Vienna (Austria): R Foundation for Statistical Computing. Available from: https://www.r-project.org/.

Robles NV, Moran BM, Rodríguez Barrera MJ, Jofre GI, Gunn T, Iverson ENK, Beskid S, Baczenas JJ, Sedghifar A, Andolfatto P, et al. 2025. Admixture mapping reveals evidence for multiple mitonuclear incompatibilities in swordtail fish hybrids. Mol Ecol. 34:e70106.

Rockenbach K, Havird JC, Monroe JG, Triant DA, Taylor DR, Sloan DB. 2016. Positive selection in rapidly evolving plastid–nuclear enzyme complexes. Genetics. 204:1507–1522.

Schwarz EN, Ruhlman TA, Sabir JSM, Khiyami MA, Hajarah NH, Alharbi NS, Al-Malki AL, Bailey CD, Jansen RK. 2017. Plastome-wide nucleotide substitution rates reveal accelerated evolution in Geraniaceae. J Mol Evol. 84:187–203.

Shi Y, Liu Y, Zhang S, Zou R, Tang J, Mu W, Peng Y, Dong S. 2018. Assembly and comparative analysis of the complete mitochondrial genome sequence of *Sophora japonica* ‘JinhuaiJ2’. PLoS One. 13:e0202485.

Skippington E, Barkman TJ, Rice DW, Palmer JD. 2015. Miniaturized mitogenome of the parasitic plant *Viscum scurruloideum* is extremely divergent and dynamic and has lost all *nad* genes. Proc Natl Acad Sci USA. 112:E3515–E3524.

Skippington E, Barkman TJ, Rice DW, Palmer JD. 2017. Comparative mitogenomics indicates respiratory competence in parasitic *Viscum* despite loss of complex I and extreme sequence divergence, and reveals horizontal gene transfer and remarkable variation in genome size. BMC Plant Biol. 17:49.

Sloan DB. 2015. Using plants to elucidate the mechanisms of cytonuclear coevolution. New Phytol. 205:1040–1046.

Sloan DB, Alverson AJ, Chuckalovcak JP, Wu M, McCauley DE, Palmer JD, Taylor DR. 2012. Rapid evolution of enormous, multichromosomal genomes in flowering plant mitochondria with exceptionally high mutation rates. PLoS Biol. 10:e1001241.

Sloan DB, Havird JC, Sharbrough J. 2017. The on-again, off-again relationship between mitochondrial genomes and species boundaries. Mol Ecol. 26:2212–2236.

Sloan DB, Warren JM, Williams AM, Wu Z, Abdel-Ghany SE, Chicco AJ, Havird JC. 2018. Cytonuclear integration and coevolution. Nat Rev Genet. 19:635–648.

Smith, C.H.; Mejia-Trujillo, R.; Havird, J.C. 2024. Mitonuclear compatibility is maintained despite relaxed selection on male mitochondrial DNA in bivalves with doubly uniparental inheritance. Evolution 78:1790–1803.

Stamatakis A, Kozlov AM, Darriba D, Flouri T, Morel B. 2019. RAxML-NG: a fast, scalable, and user-friendly tool for maximum likelihood phylogenetic inference. Bioinformatics. 35:4453–4455.

Tao M, Chen J, Cui C, Xu Y, Xu J, Shi Z, Yun J, Zhang J, Ou G-Z, Liu C, et al. 2024. Identification of a longevity gene through evolutionary rate covariation of insect mito-nuclear genomes. Nat Aging. 4:1076–1088.

Tressel LG, Shrestha B, Lee C, Choi I-S, Ruhlman TA, Cardoso D, Wojciechowski MF, Jansen RK. 2025. Plastid–nuclear coevolution of ribosomal protein genes in papilionoid legumes. Mol Phylogenet Evol. 204:108281.

Wallnoefer O, Formaggioni A, Plazzi F, Passamonti M. 2025. Convergent evolution in nuclear and mitochondrial OXPHOS subunits underlies the phylogenetic discordance in deep lineages of Squamata. Mol Phylogenet Evol. 208:108358.

Weaver RJ, Carrion G, Nix R, Maeda GP, Rabinowitz S, Iverson ENK, Thueson K, Havird JC. 2020. High mitochondrial mutation rates in Silene are associated with nuclear-mediated changes in mitochondrial physiology. Biol Lett. 16:20200450.

Weaver RJ, Rabinowitz S, Thueson L, Havird JC. 2022. Genomic signatures of mitonuclear coevolution in mammals. Mol Biol Evol. 39:msac233.

Weng M-L, Ruhlman TA, Gibby M, Jansen RK. 2012. Phylogeny, rate variation, and genome size evolution of Pelargonium (Geraniaceae). Mol Phylogenet Evol. 64:654–670.

Wertheim JO, Murrell B, Smith MD, Kosakovsky Pond SL, Scheffler K. 2015. RELAX: detecting relaxed selection in a phylogenetic framework. Mol Biol Evol. 32:820–832.

Wojciechowski MF, Lavin M, Sanderson MJ. 2004. A phylogeny of legumes (Leguminosae) based on analysis of the plastid *matK* gene resolves many well-supported subclades within the family. Am J Bot. 91:1846–1862.

Yang Z. 2007. PAML 4: phylogenetic analysis by maximum likelihood. Mol Biol Evol. 24:1586–1591.

Yan Z, Ye G, Werren JH. 2019. Evolutionary rate correlation between mitochondrial-encoded and mitochondria-associated nuclear-encoded proteins in insects. Mol Biol Evol. 36:1022–1036. doi:10.1093/molbev/msz036.

Yates AD, Allen J, Amode RM, Azov AG, Barba M, Becerra A, Bhai J, Campbell LI, Carbajo Martinez M, Chakiachvili M, et al. 2022. Ensembl Genomes 2022: an expanding genome resource for non-vertebrates. Nucleic Acids Res. 50:D996–D1003. doi:10.1093/nar/gkab1007.

Yu T, Sun L, Cui H, Liu S, Men J, Chen S, Chen Y, Lu C. 2018. The complete mitochondrial genome of a tertiary relict evergreen woody plant *Ammopiptanthus mongolicus*. Mitochondrial DNA B. 3:9–11.

Zhu A, Guo W, Jain K, Mower JP. 2014. Unprecedented heterogeneity in the synonymous substitution rate within a plant genome. Mol Biol Evol. 31:1228–1236.

