## Supplemental Figures and Tables for "Coordinated Evolutionary Rates in Oxidative Phosphorylation Complexes of Papilionoid Legumes: Cytonuclear Coevolution and Relaxed Selection"

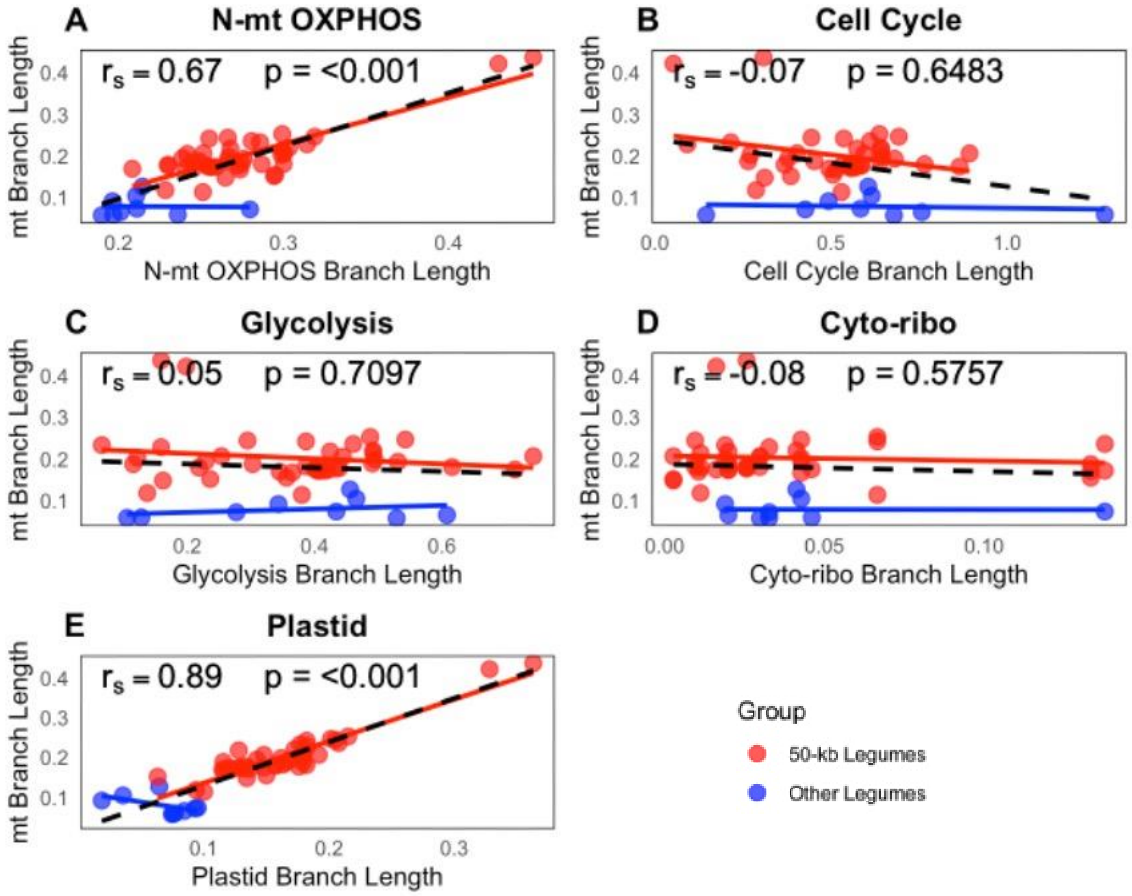

**Figure S1.** Correlation of evolutionary rates between mt and N-mt OXPHOS, cell cycle, glycolysis, cytosolic ribosomal (cyto-ribo), and plastid gene sets in the 50-kb inversion clade and other legumes based on root-to-tip branch lengths. Scatter plots display the relationship between mt branch lengths and branch lengths for each nuclear gene group. Spearman's rank correlation coefficients ($r_s$) and associated p-values are indicated for each gene group. Spearman's rank correlation coefficients ($r_s$) and associated p-values, shown on each plot, pertain to the 50-kb clade. The red regression line represents the 50-kb clade, the blue regression line corresponds to other legumes, and the black dashed line indicates the regression line for all taxa combined.

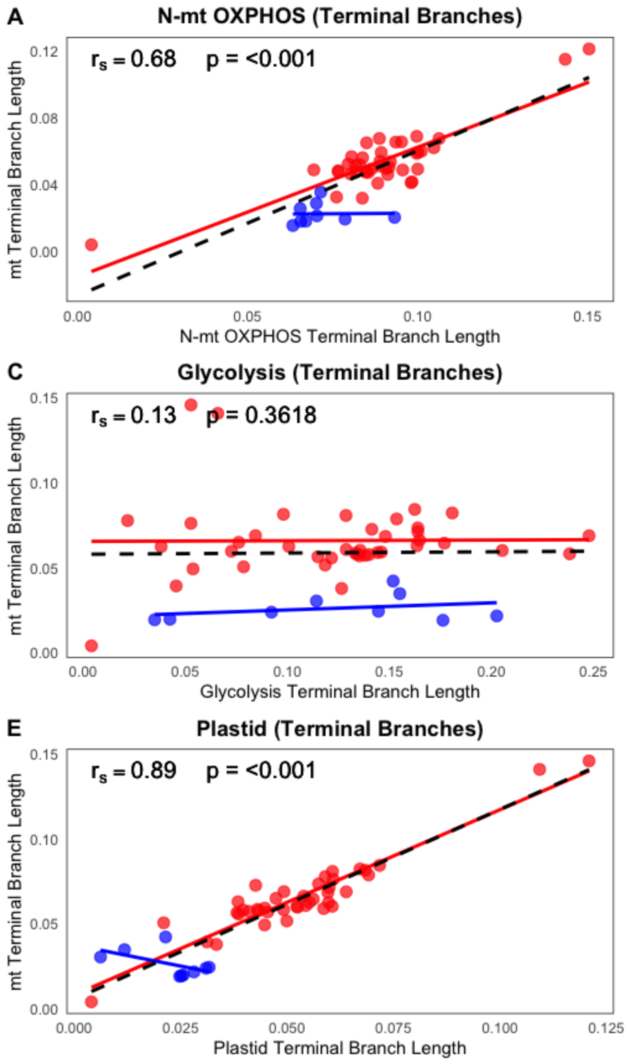

**Figure S2.** Correlation of evolutionary rates between mt and nuclear gene sets, as in Figure S1, except terminal branch lengths are used here.

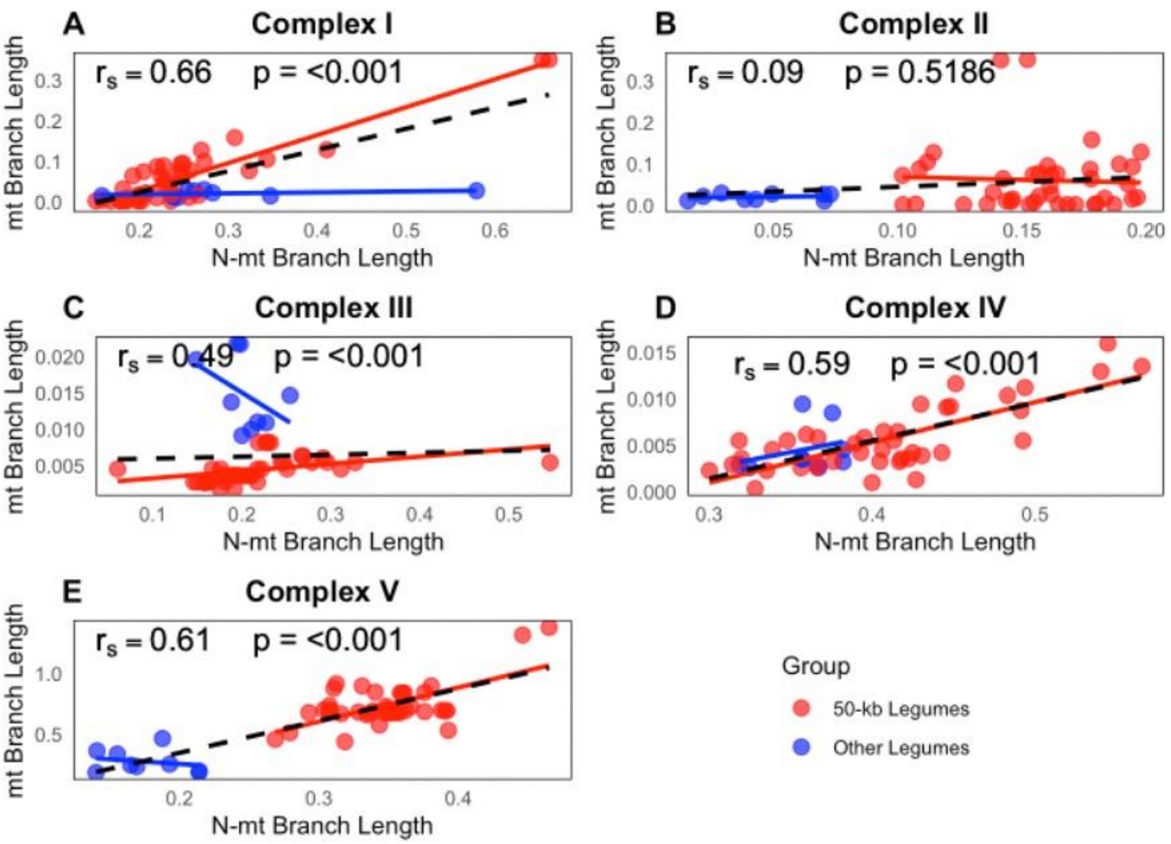

**Figure S3.** Correlation of evolutionary rates between mitochondrial-encoded and nuclear-encoded mitochondrial-targeted genes across five oxidative phosphorylation complexes in the 50-kb inversion clade and other legumes. CII has no mt-encoded genes, so the overall mt gene dataset was used for that correlation. Scatter plots display the relationship between mitochondrial branch lengths and nuclear mitochondrial-targeted branch lengths for each oxidative phosphorylation complex. Spearman's rank correlation coefficients ($r_s$) and associated p-values, shown on each plot, pertain to the 50-kb clade. The red regression line represents the 50-kb clade, the blue regression line corresponds to other legumes, and the black dashed line indicates the regression line for all taxa combined.

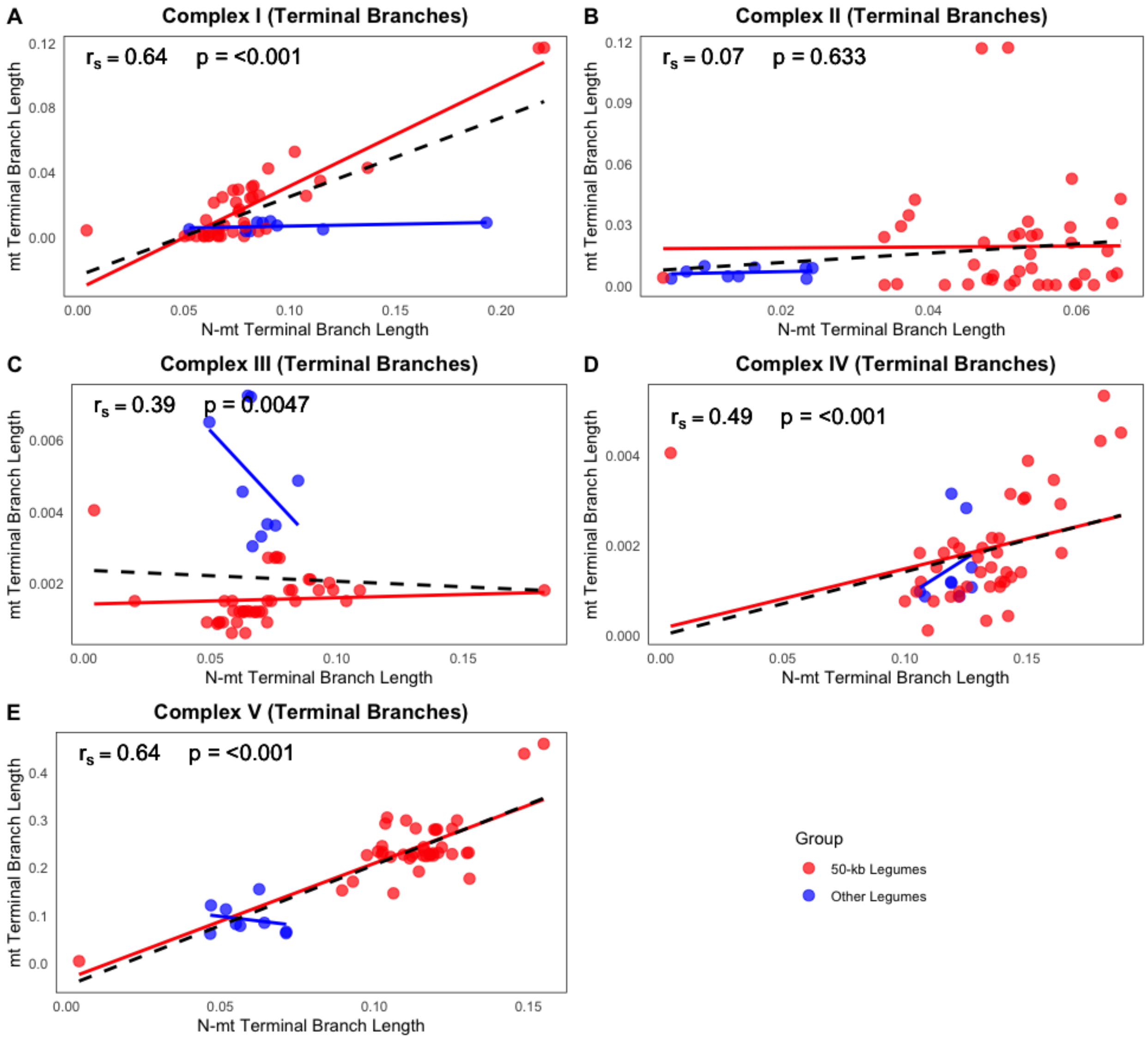

**Figure S4.** Correlation of evolutionary rates between mt and nuclear gene sets, as in Figure S3, except terminal branch lengths are used here.

**Table S1.** Fifty-nine plastid protein coding genes used for constraint tree construction.

| | | |
|---|---|---|
| *accD* | *psaB* | *rpl14* |
| *atpA* | *psaC* | *rpl23* |
| *atpB* | *psaI* | *rpl32* |
| *atpE* | *psaJ* | *rpl36* |
| *atpF* | *psbA* | *rpoA* |
| *atpH* | *psbB* | *rpoB* |
| *atpI* | *psbC* | *rpoC1* |
| *ccsA* | *psbD* | *rps2* |
| *cemA* | *psbE* | *rps3* |
| *matK* | *psbF* | *rps4* |
| *ndhC* | *psbH* | *rps7* |
| *ndhE* | *psbI* | *rps8* |
| *ndhG* | *psbJ* | *rps11* |
| *ndhI* | *psbK* | *rps12* |
| *ndhJ* | *psbL* | *rps14* |
| *petA* | *psbM* | *rps15* |
| *petG* | *psbT* | *rps18* |
| *petL* | *psbZ* | *rps19* |
| *petN* | *rbcL* | *ycf2* |
| *psaA* | *rpl2* | |

**Table S2.** Tests for relaxed selection on individual gene groups in the 50-kb inversion clade using RELAX: mt (mitochondrial), N-mt OXPHOS (nuclear-encoded mitochondrial-targeted oxidative phosphorylation), N-gly (nuclear-encoded glycolysis), N-cc (nuclear-encoded cell cycle), N-cr (nuclear-encoded cytosolic ribosomal, and N-rand (random nuclear proteins). For each gene, $p$-value indicates significance of the likelihood ratio test, and $k$ is the selection intensity parameter ($< 1$ = relaxed, $> 1$ = intensified). w values (w1–w3) correspond to purifying, neutral, and positive selection site classes, respectively, for test (50-kb inversion clade) and reference branches (other legumes), with accompanying percentages showing the proportion of sites in each class.

| Gene Group | Gene | p-value | k* | w1_test | w1_Ref | w1_% | w2_test | w2_Ref | w2_% | w3_test | w3_Ref | w3_% |
|---|---|---|---|---|---|---|---|---|---|---|---|---|
| mt | nad1 | 0.00463 | 0.65 | 0.14857 | 0.13379136 | 55.9037239 | 0.44104428 | 0.46274762 | 22.5831883 | 0.82443118 | 0.78195809 | 11.1063174 |
| mt | nad2 | 0.01856 | 0.88 | 0.07421 | 0.06793041 | 45.41664503 | 0.28461037 | 0.28935589 | 39.0810205 | 0.91075106 | 0.84059351 | 18.4645229 |
| mt | nad3 | 0.00668 | 0.793 | 0.11721 | 0.1053136 | 48.77942841 | 0.25323186 | 0.27663923 | 32.1234927 | 1.12248051 | 1.14731428 | 11.2748866 |
| mt | nad4 | 0.04329 | 0.739 | 0.12616 | 0.10585437 | 41.56912763 | 0.42518443 | 0.41454381 | 24.5728561 | 1.09930388 | 1.05283203 | 13.9728729 |
| mt | nad4L | 0.03154 | 0.562 | 0.07376 | 0.06966561 | 40.50701487 | 0.44205042 | 0.42310521 | 33.4340137 | 0.87380841 | 0.88800574 | 17.9729537 |

| | | | | | | | | | | | | |
|---|---|---|---|---|---|---|---|---|---|---|---|---|
| mt | nad5 | 0.01721 | 1.262 | 0.12282 | 0.10515289 | 59.25296829 | 0.49715154 | 0.53380147 | 32.3625648 | 0.88373973 | 0.82264905 | 11.4991743 |
| mt | nad6 | 0.00411 | 0.523 | 0.08678 | 0.06984603 | 56.71960241 | 0.3237853 | 0.30588618 | 27.1632544 | 0.94818884 | 0.94461229 | 12.292514 |
| mt | nad7 | 0.01624 | 0.846 | 0.11323 | 0.10520137 | 53.91948412 | 0.31160543 | 0.34047396 | 22.2711518 | 0.99380919 | 1.00028673 | 17.2225257 |
| mt | nad9 | 0.01693 | 0.74 | 0.11335 | 0.0946951 | 48.17905889 | 0.43292389 | 0.39068389 | 33.4314639 | 1.04730191 | 0.9534267 | 17.2003654 |
| mt | cob | 0.03675 | 0.783 | 0.10358 | 0.10234654 | 43.4658864 | 0.30224106 | 0.3306444 | 30.406154 | 0.94756546 | 0.91659983 | 16.4114763 |
| mt | cox1 | 0.03224 | 0.908 | 0.05903 | 0.05848608 | 43.12874085 | 0.4792272 | 0.43544116 | 35.4463678 | 0.98501389 | 0.91299256 | 16.9394844 |
| mt | cox3 | 0.04447 | 1.188 | 0.13353 | 0.13125637 | 45.00485796 | 0.45752383 | 0.49331528 | 30.40327 | 1.09898838 | 1.00301921 | 15.4272444 |
| mt | atp8 | 0.02414 | 0.833 | 0.08208 | 0.07174053 | 50.98453329 | 0.32869821 | 0.33051927 | 37.04363 | 0.81467328 | 0.89450478 | 12.5179906 |
| mt | atp1 | 0.00686 | 0.585 | 0.06865 | 0.05513222 | 54.29191845 | 0.42526132 | 0.46718909 | 31.0381368 | 0.90097478 | 0.86896384 | 13.4569599 |
| mt | atp6 | 0.03595 | 1.573 | 0.05408 | 0.05330469 | 53.20394753 | 0.42636286 | 0.3900194 | 31.2187594 | 1.08533983 | 1.15260365 | 11.8159772 |
| mt | atp9 | 0.03828 | 0.573 | 0.10909 | 0.09661412 | 45.59867794 | 0.23093716 | 0.23342455 | 37.5330721 | 1.15808274 | 1.10125345 | 19.0845056 |
| mt | atp4 | 0.0285 | 0.622 | 0.11776 | 0.11697465 | 59.09730561 | 0.47076587 | 0.5149522 | 28.0696573 | 1.00467098 | 1.04114108 | 15.8339179 |
| N-mt OXPHOS | NDUS1 | 0.03878 | 0.71 | 0.05166 | 0.05128412 | 54.75793833 | 0.35157571 | 0.35319984 | 22.6803046 | 1.01284539 | 1.06555951 | 14.0085142 |
| N-mt OXPHOS | NDUS4 | 0.0252 | 0.673 | 0.10121 | 0.09823462 | 51.08708105 | 0.44793724 | 0.45952973 | 20.5756535 | 0.8428688 | 0.85899099 | 14.620058 |
| N-mt OXPHOS | NDS5A | 0.02661 | 0.616 | 0.07265 | 0.06239834 | 52.23441492 | 0.29601488 | 0.30760379 | 35.1027451 | 0.97896495 | 0.97339976 | 19.4728334 |
| N-mt OXPHOS | NDS5B | 0.02195 | 0.745 | 0.11452 | 0.10043628 | 48.39200125 | 0.46865697 | 0.46439604 | 32.406191 | 1.01304691 | 0.99518505 | 11.533514 |
| N-mt OXPHOS | NDUS6 | 0.00225 | 1.556 | 0.06744 | 0.06543213 | 44.95461979 | 0.3167605 | 0.32484158 | 34.0815954 | 0.8969882 | 0.86987553 | 15.8622983 |
| N-mt OXPHOS | NDUS7 | 0.00629 | 0.617 | 0.11909 | 0.10282045 | 47.11945357 | 0.2032513 | 0.20667869 | 24.2592832 | 0.90769729 | 0.98567378 | 15.0588868 |
| N-mt OXPHOS | NDS8A | 0.00254 | 0.647 | 0.08867 | 0.07394178 | 55.15692221 | 0.47161459 | 0.50945299 | 22.7274295 | 0.95091367 | 1.01379177 | 16.1145424 |
| N-mt OXPHOS | NDS8B | 0.03218 | 0.682 | 0.14367 | 0.13093513 | 40.28786977 | 0.227386 | 0.20671418 | 20.2908933 | 0.80802848 | 0.88317948 | 10.1811018 |
| N-mt OXPHOS | NDUV1 | 0.0164 | 0.814 | 0.06375 | 0.06293597 | 42.32145281 | 0.29579409 | 0.28283613 | 27.0117512 | 0.92883167 | 0.85903874 | 18.7212391 |
| N-mt OXPHOS | NDUV2 | 0.02592 | 0.58 | 0.08411 | 0.07899661 | 40.92005284 | 0.48501859 | 0.52871018 | 31.7983537 | 0.8845792 | 0.92542332 | 19.3211828 |
| N-mt OXPHOS | NDUA1 | 0.04547 | 0.706 | 0.06135 | 0.05607465 | 40.81457605 | 0.48518214 | 0.52305195 | 27.8448809 | 0.93099894 | 1.01261784 | 15.6513318 |

| | | | | | | | | | | | | |
|---|---|---|---|---|---|---|---|---|---|---|---|---|
| N-mt OXPHOS | NDUA2 | 0.01322 | 0.737 | 0.14247 | 0.11674495 | 57.10921168 | 0.37203137 | 0.36873195 | 28.7494984 | 0.84790485 | 0.79384805 | 16.9665082 |
| N-mt OXPHOS | NDUA5 | 0.02111 | 0.519 | 0.13773 | 0.12712499 | 54.07315719 | 0.38955116 | 0.39891072 | 38.0831739 | 1.15621091 | 1.05596656 | 19.2249938 |
| N-mt OXPHOS | NDUA6 | 0.03802 | 1.743 | 0.07579 | 0.07563924 | 49.48347658 | 0.33453366 | 0.31963896 | 26.9651093 | 1.03743698 | 1.08746648 | 17.0723863 |
| N-mt OXPHOS | NDA8A | 0.01221 | 0.568 | 0.116 | 0.09604995 | 41.95668321 | 0.28796323 | 0.2700013 | 30.2797898 | 1.07164093 | 1.08760261 | 11.5253904 |
| N-mt OXPHOS | NDA8B | 0.00477 | 0.526 | 0.13172 | 0.11903088 | 49.8323175 | 0.29859936 | 0.29643144 | 35.6730603 | 1.1156685 | 1.19194201 | 15.7628836 |
| N-mt OXPHOS | NDUA9 | 0.0152 | 0.88 | 0.10552 | 0.10293208 | 49.46943542 | 0.40175554 | 0.38997223 | 27.9308556 | 0.99937688 | 0.92737625 | 16.0671505 |
| N-mt OXPHOS | NDUAC | 0.0089 | 0.886 | 0.10297 | 0.09763139 | 43.4640374 | 0.42571236 | 0.43283505 | 32.441734 | 0.83476812 | 0.88406406 | 14.2413067 |
| N-mt OXPHOS | NDADB | 0.04656 | 1.823 | 0.07419 | 0.06969432 | 48.67703298 | 0.43747371 | 0.40052771 | 37.2472742 | 1.01484262 | 0.95428235 | 17.3644424 |
| N-mt OXPHOS | NDADA | 0.0406 | 0.622 | 0.05931 | 0.05578087 | 47.97009469 | 0.43688544 | 0.47833668 | 38.9904125 | 1.03473645 | 0.96513096 | 19.3436701 |
| N-mt OXPHOS | NDUB7 | 0.03204 | 0.539 | 0.13972 | 0.12182162 | 52.31700196 | 0.22736183 | 0.24947098 | 22.9414696 | 1.09817579 | 1.02443676 | 19.2556851 |
| N-mt OXPHOS | NDUB9 | 0.0437 | 1.374 | 0.14004 | 0.12025492 | 52.70187302 | 0.34832609 | 0.36213107 | 38.5317525 | 0.97266382 | 1.03385891 | 14.5083937 |
| N-mt OXPHOS | NDBAA | 0.04038 | 0.676 | 0.11331 | 0.10898974 | 40.9060802 | 0.21726763 | 0.21883614 | 29.8423259 | 0.85103212 | 0.87914975 | 11.1323805 |
| N-mt OXPHOS | NDB3 | 0.01014 | 0.549 | 0.0839 | 0.0807137 | 47.49225229 | 0.36485866 | 0.35095956 | 25.1648878 | 0.91351036 | 0.91772446 | 19.848412 |
| N-mt OXPHOS | NDB4 | 0.04474 | 0.698 | 0.08492 | 0.08266236 | 52.51719831 | 0.33245915 | 0.35332396 | 29.1827151 | 0.94523292 | 0.9185453 | 18.3889809 |
| N-mt OXPHOS | NDA2 | 0.02743 | 0.514 | 0.1226 | 0.12047266 | 50.06272517 | 0.46631125 | 0.4835397 | 39.6006515 | 1.0583669 | 1.13821021 | 11.2466268 |
| N-mt OXPHOS | NDB1 | 0.04056 | 0.864 | 0.13971 | 0.12605593 | 57.12979682 | 0.3052745 | 0.28467561 | 29.8523619 | 1.02831132 | 1.00619134 | 19.2084188 |
| N-mt OXPHOS | NDA1 | 0.04491 | 0.604 | 0.13871 | 0.12488107 | 53.17387263 | 0.2351201 | 0.25444355 | 26.5750322 | 0.94243869 | 1.0021138 | 18.6989636 |
| N-mt OXPHOS | NDB2 | 0.01658 | 0.765 | 0.12799 | 0.12282676 | 43.25868854 | 0.2428975 | 0.2585662 | 32.6680171 | 1.1946061 | 1.18006414 | 15.1883806 |
| N-mt OXPHOS | SDHA1 | 0.00639 | 0.625 | 0.1142 | 0.10620518 | 41.41137495 | 0.42845319 | 0.46699683 | 24.8029124 | 1.04230993 | 1.01665752 | 15.9127544 |
| N-mt OXPHOS | SDHA2 | 0.01217 | 0.708 | 0.05841 | 0.05492838 | 52.84838556 | 0.38546542 | 0.40286683 | 21.5172666 | 0.89489072 | 0.88821121 | 13.990027 |
| N-mt OXPHOS | SDHB2 | 0.02193 | 0.719 | 0.06616 | 0.06345793 | 40.53022621 | 0.2303368 | 0.23556154 | 22.5775944 | 0.84071299 | 0.80731615 | 10.5476164 |
| N-mt OXPHOS | SDHB3 | 0.04108 | 0.574 | 0.13986 | 0.13678323 | 51.71551163 | 0.22523204 | 0.22154918 | 22.5609168 | 0.86114366 | 0.90378911 | 13.3519724 |
| N-mt OXPHOS | SDHB1 | 0.04318 | 0.888 | 0.11064 | 0.09599116 | 58.80460483 | 0.41029074 | 0.44579964 | 23.0380539 | 0.89838309 | 0.89887188 | 18.0285345 |

| | | | | | | | | | | | | |
|---|---|---|---|---|---|---|---|---|---|---|---|---|
| N-mt OXPHOS | SDH31 | 0.00134 | 1.81 | 0.05092 | 0.04456094 | 51.50948356 | 0.2218289 | 0.23806961 | 22.7765435 | 0.86427255 | 0.81798431 | 10.0463202 |
| N-mt OXPHOS | SDH4 | 0.02603 | 0.876 | 0.06015 | 0.0492506 | 47.76339852 | 0.44655802 | 0.40594077 | 32.8174949 | 0.87462681 | 0.94452254 | 13.3349917 |
| N-mt OXPHOS | UCRI1 | 0.02145 | 0.858 | 0.11635 | 0.10653658 | 52.86576437 | 0.41187267 | 0.37285737 | 23.6376017 | 0.91403807 | 0.8928125 | 13.9816869 |
| N-mt OXPHOS | UCRI2 | 0.01188 | 0.739 | 0.05051 | 0.04077109 | 49.16505781 | 0.22440463 | 0.2188602 | 26.9133457 | 0.86934944 | 0.87692197 | 15.373956 |
| N-mt OXPHOS | CYB | 0.00687 | 0.869 | 0.06608 | 0.05901734 | 50.91233579 | 0.22545131 | 0.23945425 | 37.9357682 | 1.15870617 | 1.25290252 | 19.1985562 |
| N-mt OXPHOS | CYC1B | 0.01754 | 0.535 | 0.10487 | 0.09527743 | 58.82929618 | 0.49599187 | 0.54432887 | 29.4792328 | 0.8320935 | 0.85276902 | 13.4634599 |
| N-mt OXPHOS | QCR61 | 0.0472 | 0.578 | 0.11919 | 0.10218257 | 47.72205276 | 0.31228124 | 0.29044759 | 33.3511548 | 1.00980456 | 0.93243294 | 13.469532 |
| N-mt OXPHOS | QCR62 | 0.01684 | 0.518 | 0.1152 | 0.1057728 | 59.22381128 | 0.31119264 | 0.3170512 | 23.4463974 | 0.96415873 | 1.04897233 | 17.3750125 |
| N-mt OXPHOS | QCR71 | 0.02642 | 1.33 | 0.07243 | 0.05838583 | 58.10701284 | 0.44383987 | 0.43326679 | 23.8457804 | 1.19295145 | 1.22342135 | 14.5221794 |
| N-mt OXPHOS | QCR72 | 0.03545 | 0.655 | 0.12122 | 0.09788147 | 43.9158227 | 0.48417457 | 0.52967869 | 20.8173723 | 0.84481556 | 0.8169207 | 12.2460482 |
| N-mt OXPHOS | QCR9 | 0.01882 | 0.609 | 0.07372 | 0.07110442 | 41.38722602 | 0.49580032 | 0.52972485 | 23.3787013 | 0.95914224 | 0.88994436 | 14.5243952 |
| N-mt OXPHOS | COX11 | 0.04862 | 1.831 | 0.08254 | 0.07197803 | 42.01556003 | 0.42601346 | 0.45483997 | 25.5718068 | 1.18778817 | 1.2576361 | 11.4085702 |
| N-mt OXPHOS | COX15 | 0.04816 | 0.643 | 0.12465 | 0.10288762 | 40.36443651 | 0.31287788 | 0.31091883 | 23.5402097 | 1.14620285 | 1.1737284 | 11.7638699 |
| N-mt OXPHOS | CX171 | 0.01334 | 0.612 | 0.11496 | 0.10397542 | 41.88885922 | 0.22505022 | 0.22121624 | 21.7740507 | 1.12682883 | 1.13436981 | 14.9836777 |
| N-mt OXPHOS | CX172 | 0.02537 | 0.717 | 0.13492 | 0.12871351 | 53.66013547 | 0.43314407 | 0.4135146 | 22.4127174 | 0.90316113 | 0.97431082 | 14.1892545 |
| N-mt OXPHOS | CYC2 | 0.01574 | 0.556 | 0.11576 | 0.09760469 | 41.42377297 | 0.36752127 | 0.33491299 | 29.2155754 | 0.86835503 | 0.918476 | 19.148459 |
| N-mt OXPHOS | CYC1 | 0.01496 | 0.821 | 0.10683 | 0.09877268 | 46.37951261 | 0.3272666 | 0.35113889 | 24.1266744 | 1.06745729 | 0.99309285 | 13.623939 |
| N-mt OXPHOS | ATPBM | 0.00281 | 0.53 | 0.05937 | 0.04850942 | 56.89750622 | 0.47190632 | 0.50143831 | 27.2853972 | 1.1717504 | 1.12762745 | 15.8058835 |
| N-mt OXPHOS | ATPBO | 0.03087 | 0.895 | 0.08677 | 0.07031288 | 40.46543871 | 0.23335924 | 0.25668199 | 30.0683454 | 1.02270516 | 0.97126087 | 16.3226429 |
| N-mt OXPHOS | ATPG3 | 0.02563 | 0.809 | 0.07652 | 0.06934785 | 56.28936965 | 0.34778753 | 0.38233235 | 33.8078966 | 1.02864508 | 1.07883191 | 10.1309446 |
| N-mt OXPHOS | ATP4 | 0.00352 | 0.579 | 0.0744 | 0.06756465 | 45.6370955 | 0.20340609 | 0.20566112 | 20.7862428 | 0.91199164 | 0.82690873 | 16.6353737 |
| N-mt OXPHOS | ATP5E | 0.01465 | 0.502 | 0.1473 | 0.13661868 | 42.36329655 | 0.34059819 | 0.35892152 | 35.988208 | 1.10779717 | 1.12328189 | 11.7803597 |
| N-mt OXPHOS | ATPO | 0.04551 | 0.826 | 0.08931 | 0.08441744 | 53.93474331 | 0.21689098 | 0.23618412 | 32.5580078 | 0.8748175 | 0.92073819 | 19.6107032 |

| | | | | | | | | | | | | |
|---|---|---|---|---|---|---|---|---|---|---|---|---|
| N-mt OXPHOS | ATP61 | 0.01274 | 0.783 | 0.1392 | 0.13852772 | 52.57885694 | 0.23564537 | 0.25212393 | 21.6351806 | 0.92947169 | 0.99951029 | 11.4866273 |
| N-mt OXPHOS | ATP9 | 0.0081 | 0.792 | 0.11311 | 0.10216775 | 57.54944027 | 0.23525787 | 0.2233702 | 37.4715725 | 0.97017458 | 0.93953292 | 14.1462412 |
| N-mt OXPHOS | ATP5H | 0.02498 | 0.809 | 0.12948 | 0.11194728 | 54.70142088 | 0.39476309 | 0.39085842 | 38.417448 | 1.00304415 | 1.0674912 | 10.8534967 |
| N-gly | G3PC2 | 0.36584889 | 1.04996448 | 0.09220195 | 0.13155455 | 0.212169768 | 0.08183132 | 0.22497775 | 0.63698432 | 0.17617773 | 0.19058577 | 0.15084591 |
| N-gly | PDK | 0.5881374 | 1.01283231 | 0.16847241 | 0.06009436 | 0.781398111 | 0.12552593 | 0.11198435 | 0.2182478 | 0.1472845 | 0.10287344 | 0.00035409 |
| N-gly | PFKA3 | 0.80277887 | 1.04322057 | 0.09717547 | 0.13583128 | 0.349791403 | 0.06052636 | 0.24351676 | 0.48152697 | 0.08994162 | 0.18104063 | 0.16868162 |
| N-gly | ALFC6 | 0.69469518 | 1.04008111 | 0.14288967 | 0.0943406 | 0.731537404 | 0.11140619 | 0.16711163 | 0.26470625 | 0.13242387 | 0.20425673 | 0.00375634 |
| N-gly | TPIS | 0.33622147 | 1.05 | 0.05180973 | 0.03110334 | 0.637347747 | 0.228254 | 0.03600237 | 0.31854413 | 0.20555302 | 0.20346595 | 0.04410812 |
| N-gly | 6PGL1 | 0.35072833 | 0.99244627 | 0.17478659 | 0.04223685 | 0.694649981 | 0.15607364 | 0.12209006 | 0.24683983 | 0.23360816 | 0.15785826 | 0.05851019 |
| N-gly | G6PI | 0.40043857 | 0.99964432 | 0.16677699 | 0.13616582 | 0.959783544 | 0.13467796 | 0.19562965 | 0.03559851 | 0.08197098 | 0.17386152 | 0.00461795 |
| N-gly | PMG1 | 0.79511767 | 1.04351093 | 0.03587133 | 0.06973007 | 0.511511585 | 0.11837036 | 0.10597171 | 0.2105106 | 0.10019773 | 0.0746286 | 0.27797781 |
| N-gly | DPNP3 | 0.89338639 | 1.02333452 | 0.18493713 | 0.16739729 | 0.622160627 | 0.2401612 | 0.12121792 | 0.18746623 | 0.2298452 | 0.23330775 | 0.19037314 |
| N-gly | HXK2 | 0.54694026 | 1.00621832 | 0.07030686 | 0.06715688 | 0.858736985 | 0.18576115 | 0.1173331 | 0.07464322 | 0.24725287 | 0.22872898 | 0.06661979 |
| N-cc | RBR1 | 0.89331683 | 1.00714365 | 0.04756377 | 0.07149699 | 0.551061259 | 0.15866712 | 0.15769021 | 0.19267957 | 0.23025666 | 0.21638947 | 0.25625917 |
| N-cc | E2FB | 0.81277596 | 0.99737495 | 0.06453006 | 0.09967683 | 0.610422222 | 0.17959677 | 0.20113351 | 0.18448674 | 0.23945776 | 0.19220409 | 0.20509104 |
| N-cc | CKS1 | 0.39485168 | 0.99132383 | 0.16997848 | 0.10443128 | 0.706985133 | 0.21330986 | 0.17039197 | 0.24885408 | 0.09890374 | 0.24644836 | 0.04416079 |
| N-cc | CDC2C | 0.74804548 | 0.98581698 | 0.14955255 | 0.14690257 | 0.557740779 | 0.20147347 | 0.21070978 | 0.37528139 | 0.08952853 | 0.20045199 | 0.06697783 |
| N-cc | DPA | 0.66263977 | 1.05 | 0.16462358 | 0.17589739 | 0.557266093 | 0.24255819 | 0.11410075 | 0.43718049 | 0.17365468 | 0.16369721 | 0.00555342 |
| N-cc | KRP1 | 0.63192497 | 1.01068791 | 0.13885481 | 0.16451624 | 0.866695589 | 0.16950656 | 0.0849223 | 0.12892365 | 0.06678016 | 0.16618587 | 0.00438076 |
| N-cr | RPL10AA | 0.69665596 | 0.99381521 | 0.13574494 | 0.04335497 | 0.640376565 | 0.21043426 | 0.24611684 | 0.3584571 | 0.05328895 | 0.14208317 | 0.00116633 |
| N-cr | RPL12A | 0.88887083 | 1.00892014 | 0.14459579 | 0.14468138 | 0.882471016 | 0.08404758 | 0.15005014 | 0.06986174 | 0.2193962 | 0.20037629 | 0.04766725 |
| N-cr | RPL17A | 0.41434819 | 1.00818795 | 0.15538966 | 0.14317965 | 0.956847037 | 0.11631187 | 0.23897396 | 0.03816867 | 0.09080039 | 0.24838562 | 0.00498429 |
| N-cr | RPL23AA | 0.86672099 | 0.9912692 | 0.02093756 | 0.17840699 | 0.45732142 | 0.21520194 | 0.18740073 | 0.54014951 | 0.12115115 | 0.23186185 | 0.00252907 |

| | | | | | | | | | | | | |
|---|---|---|---|---|---|---|---|---|---|---|---|---|
| N-cr | RPL26B | 0.50391255 | 0.99111161 | 0.18964668 | 0.09377828 | 0.719148081 | 0.08202797 | 0.11012002 | 0.15677982 | 0.23061425 | 0.09645571 | 0.1240721 |
| N-cr | RPL3A | 0.7643906 | 1.02479431 | 0.05338956 | 0.02502734 | 0.733589503 | 0.11370959 | 0.11308364 | 0.1440359 | 0.22128442 | 0.08727837 | 0.12237459 |
| N-cr | RPL3B | 0.46785621 | 1.02642805 | 0.1793081 | 0.08594658 | 0.711596198 | 0.06382206 | 0.05509464 | 0.28531561 | 0.20882268 | 0.21957634 | 0.00308819 |
| N-cr | RPL4A | 0.44845818 | 1.05 | 0.07181619 | 0.02742042 | 0.509861942 | 0.07402236 | 0.0440838 | 0.48728716 | 0.10598724 | 0.23444512 | 0.00285089 |
| N-cr | RPL7A | 0.33719123 | 0.99679427 | 0.1105243 | 0.17887903 | 0.588239752 | 0.05059153 | 0.1998933 | 0.19616316 | 0.15243718 | 0.20804955 | 0.21559709 |
| N-cr | RPL8B | 0.7065914 | 1.02336264 | 0.07153257 | 0.16531692 | 0.908900429 | 0.24900381 | 0.23221423 | 0.03715483 | 0.07987864 | 0.22491903 | 0.05394474 |
| N-cr | RPP0C | 0.51903664 | 1.00305765 | 0.0393084 | 0.10120638 | 0.693608657 | 0.17646863 | 0.06607723 | 0.21204264 | 0.18749887 | 0.12222877 | 0.0943487 |
| N-cr | RPS13B | 0.55773383 | 1.01200937 | 0.19840992 | 0.15313166 | 0.762870377 | 0.24560121 | 0.08091252 | 0.19362279 | 0.17542731 | 0.19910456 | 0.04350683 |
| N-cr | RPS15AE | 0.89410855 | 1.03920173 | 0.05583411 | 0.19902796 | 0.865382389 | 0.09552179 | 0.07100326 | 0.07306844 | 0.13118367 | 0.10673042 | 0.06154917 |
| N-cr | RPS20C | 0.48917231 | 1.04259146 | 0.05177754 | 0.06138248 | 0.297899089 | 0.14701294 | 0.08871305 | 0.6958984 | 0.22176677 | 0.08402876 | 0.00620251 |
| N-cr | RPS5B | 0.79421824 | 1.02951049 | 0.12624815 | 0.17655848 | 0.551846266 | 0.20892666 | 0.23128551 | 0.37260429 | 0.19239431 | 0.20444023 | 0.07554944 |
| N-cr | RPS9C | 0.63155299 | 1.03696496 | 0.06564646 | 0.07763886 | 0.850525867 | 0.08035348 | 0.15722351 | 0.14242251 | 0.22982286 | 0.24802067 | 0.00705162 |
| N-cr | RPSaA | 0.375028 | 1.00061187 | 0.06775117 | 0.10263843 | 0.617761322 | 0.20431108 | 0.10800442 | 0.32578171 | 0.1797109 | 0.16751285 | 0.05645697 |
| N-rand | MDN1 | 0.42462954 | 0.97661259 | 0.09650128 | 0.09247385 | 0.77975828 | 0.32124961 | 0.28136472 | 0.19857839 | 1.13493672 | 1.09785610 | 0.02323746 |
| N-rand | REV1 | 0.64715563 | 1.02162134 | 0.08864320 | 0.06674239 | 0.66865342 | 0.31179854 | 0.28576843 | 0.31222232 | 0.83998123 | 0.79815824 | 0.02101282 |
| N-rand | 1MMP | 0.65559242 | 1.00983490 | 0.13137128 | 0.12774034 | 0.90649022 | 0.33238902 | 0.37751923 | 0.08363286 | 0.94675213 | 1.09189389 | 0.01590279 |
| N-rand | SRS2 | 0.83333343 | 0.99754912 | 0.09917348 | 0.09261093 | 0.80558234 | 0.39692731 | 0.42722222 | 0.13948321 | 1.20463909 | 1.16473290 | 0.05667390 |
| N-rand | IDM1 | 0.36625434 | 0.97184731 | 0.10641111 | 0.11164581 | 0.66613244 | 0.23883267 | 0.24625412 | 0.26364233 | 1.37825269 | 1.31998563 | 0.07112987 |
| N-rand | CERK1 | 0.59846756 | 1.03584600 | 0.09845322 | 0.19740098 | 0.72435732 | 0.38912111 | 0.36686456 | 0.25698683 | 1.02133333 | 0.83860923 | 0.02476434 |
| N-rand | LNK1 | 0.47484567 | 1.00555549 | 0.07592489 | 0.06945721 | 0.84995637 | 0.26894738 | 0.28373976 | 0.12656385 | 1.12949023 | 1.15232987 | 0.02440938 |
| N-rand | RPP1 | 0.59248390 | 1.02538490 | 0.14895738 | 0.14887212 | 0.63947839 | 0.35546375 | 0.36999999 | 0.22029873 | 1.11657390 | 1.34893094 | 0.15563889 |
| N-rand | SAM2 | 0.86995783 | 1.03438948 | 0.10551487 | 0.11991093 | 0.66479039 | 0.25693748 | 0.26647294 | 0.29197382 | 1.01347289 | 1.19174563 | 0.04555555 |
| N-rand | BCB | 0.85663902 | 1.02685673 | 0.14782838 | 0.15375839 | 0.80938432 | 0.24967482 | 0.29373823 | 0.08859632 | 1.16943245 | 1.04926474 | 0.10335833 |

| | | | | | | | | | | | | |
|---|---|---|---|---|---|---|---|---|---|---|---|---|
| N-rand | BZIP34 | 0.75648394 | 1.01657580 | 0.07370454 | 0.05552111 | 0.85553876 | 0.21292847 | 0.19228578 | 0.10970980 | 1.153 | 1.22365121 | 0.03663434 |
| N-rand | TK1B | 0.76984343 | 1.03384935 | 0.14246887 | 0.12373978 | 0.62352864 | 0.33695257 | 0.33183954 | 0.33434567 | 0.91564897 | 1.03901368 | 0.04457438 |
| N-rand | SAHH1 | 0.46775479 | 1.03428088 | 0.08285356 | 0.06974268 | 0.80675369 | 0.27386423 | 0.29694268 | 0.19032589 | 0.86000097 | 0.87585467 | 0.00385343 |
| N-rand | CYP98A3 | 0.61195479 | 1.03209468 | 0.08384259 | 0.09686380 | 0.55746789 | 0.28649649 | 0.29229859 | 0.44705285 | 1.03693699 | 1.11635742 | 0.00323457 |
| N-rand | KNL2 | 0.78396369 | 0.98785379 | 0.12175369 | 0.14175280 | 0.63364268 | 0.39894357 | 0.38475436 | 0.15685324 | 0.99284246 | 0.83235686 | 0.21845267 |
| N-rand | AGL15 | 0.76245806 | 1.01297647 | 0.12595367 | 0.12223235 | 0.92495625 | 0.26347694 | 0.29446769 | 0.02583950 | 1.10446284 | 1.02536395 | 0.05146285 |
| N-rand | GL2 | 0.64437289 | 1.00636294 | 0.14735628 | 0.16463840 | 0.73926743 | 0.36557394 | 0.36177777 | 0.17754738 | 0.92945375 | 1.00645639 | 0.09256389 |
| N-rand | APY6 | 0.81174954 | 1.06847303 | 0.14657638 | 0.16464842 | 0.79585635 | 0.31116372 | 0.31213620 | 0.16326483 | 1.39863265 | 1.38828563 | 0.04236484 |
| N-rand | HUB1 | 0.81193738 | 1.00783637 | 0.14336583 | 0.12993733 | 0.88663483 | 0.22536489 | 0.24227283 | 0.05536287 | 0.99692728 | 1.07946382 | 0.05999993 |
| N-rand | GRF1 | 0.52334738 | 1.04545745 | 0.11447398 | 0.12446474 | 0.34637282 | 0.24937393 | 0.22598475 | 0.64836387 | 1.17136485 | 1.15463833 | 0.00637575 |

**Table S3.** Tests for relaxed selection on concatenated gene datasets (as in Table S2).

| Gene Group | p-value | k* | w1_test | w1_Ref | w1_% | w2_test | w2_Ref | w2_% | w3_test | w3_Ref | w3_% |
|---|---|---|---|---|---|---|---|---|---|---|---|
| mt | 0.001 | 0.83 | 0.05 | 0.03 | 0.7 | 0.45 | 0.3 | 0.25 | 2.1 | 1.3 | 0.05 |
| N-mt OXPHOS | 0.002 | 0.8 | 0.04 | 0.03 | 0.72 | 0.5 | 0.35 | 0.23 | 1.9 | 1.2 | 0.05 |
| N-gly | 0.46 | 1.03 | 0.08 | 0.07 | 0.68 | 0.35 | 0.33 | 0.27 | 1.2 | 1.1 | 0.05 |
| N-cc | 0.53 | 1.01 | 0.07 | 0.06 | 0.7 | 0.3 | 0.28 | 0.25 | 1 | 0.95 | 0.05 |
| N-cr | 0.48 | 1.02 | 0.06 | 0.06 | 0.73 | 0.32 | 0.3 | 0.22 | 1.1 | 1 | 0.05 |
| N-rand | 0.41 | 1.01 | 0.05 | 0.05 | 0.69 | 0.33 | 0.27 | 0.24 | 1.2 | 0.94 | 0.05 |

**Table S4.** Summary of evolutionary rate correlation analyses across mitochondrial and nuclear gene groups in the 50-kb inversion clade. Spearman's rank correlation coefficients ($r_S$) are shown for comparisons between mt genes, N-mt OXPHOS genes, and other nuclear gene groups (glycolysis, cell cycle, and cytosolic ribosomal). ERC results are further detailed for individual OXPHOS complexes (I–V), including contact and non-contact protein categories within mitonuclear complexes.

| Comparison | Correlation Coefficient ($r_S$) |
|---|---|
| mt and N-mt OXPHOS | 0.67 ($P$ < 0.001, 95% CI = 0.44-0.83) |
| mt and Glycolysis | 0.05 ($P$ = 0.710, 95% CI = -0.26-0.35) |
| mt and Cell cycle | -0.07 ($P$ = 0.648, 95% CI = -0.37-0.25) |
| mt and Cytosolic ribosomal | -0.08 ($P$ = 0.576, 95% CI = -0.35-0.21) |
| mt and plastid | 0.89 ($P$ < 0.001, 95% CI = 0.78-0.94) |
| mt and N-mt OXPHOS (Complex I) | 0.66 ($P$ < 0.001, 95% CI = 0.44-0.81) |
| mt and N-mt OXPHOS (Complex II) | 0.09 ($P$ = 0.519, 95% CI = -0.17-0.36) |
| mt and N-mt OXPHOS (Complex III) | 0.49 ($P$ < 0.001, 95% CI = 0.20-0.67) |
| mt and N-mt OXPHOS (Complex IV) | 0.59 ($P$ < 0.001, 95% CI = 0.33-0.77) |
| mt and N-mt OXPHOS (Complex V) | 0.61 ($P$ < 0.001, 95% CI = 0.33-0.78) |
| Contact | 0.51 (95% CI = 0.23-0.73) |
| Non-contact | 0.41 (95% CI = 0.11-0.67) |
| Difference between Contact and Non-contact | $\Delta r_S$ = 0.10 ($P$ = 0.5072) |

**Table S5.** Species-level synonymous substitution rates ($d_S$) estimated from concatenated alignments of mt, N-mt OXPHOS, Glycolysis, Cell Cycle, Cytosolic ribosomal, and Random nuclear groups.

| Species | Group | mt $d_S$ | N-mt OXPHOS $d_S$ | Glycolysis $d_S$ | Cell Cycle $d_S$ | Cytosolic ribosomal $d_S$ | Random nuclear $d_S$ |
|---|---|---|---|---|---|---|---|
| *Cercis canadensis* | Other legumes | 0.0672 | 0.1686 | 0.185 | 0.1656 | 0.19 | 0.1643 |
| *Parkinsonia aculeata* | Other legumes | 0.1027 | 0.185 | 0.1604 | 0.18 | 0.1703 | 0.154 |
| *Andira humilis* | 50-kb inversion clade | 0.1209 | 0.2016 | 0.275 | 0.2038 | 0.1611 | 0.1619 |
| *Hymenolobium janeirense* | 50-kb inversion clade | 0.1777 | 0.208 | 0.2131 | 0.184 | 0.1932 | 0.168 |
| *Exostyles venusta* | 50-kb inversion clade | 0.04 | 0.2069 | 0.2447 | 0.2026 | 0.1619 | 0.1983 |
| *Harleyodendron unifoliolatum* | 50-kb inversion clade | 0.2422 | 0.1804 | 0.2438 | 0.211 | 0.1768 | 0.166 |
| *Holocalyx balansae* | 50-kb inversion clade | 0.2006 | 0.1905 | 0.2142 | 0.2193 | 0.177 | 0.1872 |
| *Zollernia ilicifolia* | 50-kb inversion clade | 0.1505 | 0.1967 | 0.2548 | 0.2036 | 0.1539 | 0.1629 |
| *Luetzelburgia bahiensis* | 50-kb inversion clade | 0.1473 | 0.1856 | 0.2366 | 0.2002 | 0.1513 | 0.1768 |
| *Sweetia fruticosa* | 50-kb inversion clade | 0.1835 | 0.2094 | 0.2409 | 0.2057 | 0.1632 | 0.1794 |
| *Vatairea guianensis* | 50-kb inversion clade | 0.1525 | 0.2086 | 0.2171 | 0.2077 | 0.1806 | 0.177 |
| *Dermatophyllum secundiflorum* | 50-kb inversion clade | 0.1537 | 0.1693 | 0.2535 | 0.1866 | 0.1617 | 0.1819 |
| *Bowdichia virgilioides* | 50-kb inversion clade | 0.2052 | 0.1701 | 0.2145 | 0.2204 | 0.1608 | 0.1606 |
| *Diplotropis ferruginea* | 50-kb inversion clade | 0.1928 | 0.1792 | 0.1975 | 0.24 | 0.146 | 0.1897 |
| *Camoensia scandens* | 50-kb inversion clade | 0.1622 | 0.2034 | 0.2552 | 0.2182 | 0.1633 | 0.1824 |
| *Baptisia leucophaea* | 50-kb inversion clade | 0.16 | 0.235 | 0.2212 | 0.1819 | 0.1731 | 0.1601 |

| | | | | | | | |
|---|---|---|---|---|---|---|---|
| *Cyclolobium brasiliense* | 50-kb inversion clade | 0.1738 | 0.1951 | 0.23 | 0.2058 | 0.1767 | 0.1781 |
| *Harpalyce brasiliana* | 50-kb inversion clade | 0.131 | 0.2033 | 0.2635 | 0.1775 | 0.1765 | 0.1551 |
| *Poecilanthe parviflora* | 50-kb inversion clade | 0.143 | 0.1813 | 0.25 | 0.1769 | 0.1653 | 0.1697 |
| *Tabaroa caatingicola* | 50-kb inversion clade | 0.1961 | 0.1843 | 0.241 | 0.2156 | 0.1662 | 0.1526 |
| *Ormosia bahiensis* | 50-kb inversion clade | 0.1574 | 0.1677 | 0.2489 | 0.2074 | 0.1823 | 0.1801 |
| *Acosmium lentiscifolium* | 50-kb inversion clade | 0.0887 | 0.1996 | 0.2172 | 0.1939 | 0.1684 | 0.205 |
| *Centrolobium tomentosum* | 50-kb inversion clade | 0.138 | 0.2153 | 0.2434 | 0.175 | 0.145 | 0.1907 |
| *Dalbergia nigra* | 50-kb inversion clade | 0.2004 | 0.1936 | 0.2495 | 0.1842 | 0.1862 | 0.1803 |
| *Platymiscium pubescens* | 50-kb inversion clade | 0.1512 | 0.2234 | 0.2431 | 0.2069 | 0.1523 | 0.1581 |
| *Poiretia bahiana* | 50-kb inversion clade | 0.1562 | 0.2043 | 0.2255 | 0.2053 | 0.1895 | 0.1762 |
| *Stylosanthes scabra* | 50-kb inversion clade | 0.2012 | 0.1998 | 0.19 | 0.2165 | 0.1964 | 0.1861 |
| *Zornia myriadena* | 50-kb inversion clade | 0.28 | 0.2083 | 0.2399 | 0.2287 | 0.1819 | 0.1691 |
| *Dalea aurea* | 50-kb inversion clade | 0.126 | 0.2079 | 0.2332 | 0.1995 | 0.1841 | 0.1711 |
| *Eysenhardia texana* | 50-kb inversion clade | 0.2383 | 0.2176 | 0.2244 | 0.2126 | 0.1977 | 0.1455 |
| *Alhagi sparsifolia* | 50-kb inversion clade | 0.0975 | 0.187 | 0.245 | 0.2023 | 0.1531 | 0.1584 |
| *Trifolium repens* | 50-kb inversion clade | 0.1734 | 0.165 | 0.2362 | 0.1964 | 0.1739 | 0.145 |
| *Coursetia rostrata* | 50-kb inversion clade | 0.1002 | 0.2012 | 0.2328 | 0.2011 | 0.1729 | 0.1969 |
| *Clitoria mariana* | 50-kb inversion clade | 0.2409 | 0.1908 | 0.2038 | 0.221 | 0.1544 | 0.1768 |
| *Periandra mediterranea* | 50-kb inversion clade | 0.2105 | 0.1986 | 0.2043 | 0.1881 | 0.205 | 0.1807 |
| *Macroptilium erythroloma* | 50-kb inversion clade | 0.2284 | 0.2263 | 0.237 | 0.2227 | 0.1614 | 0.1664 |
| *Phaseolus acutifolius* | 50-kb inversion clade | 0.1166 | 0.1808 | 0.2329 | 0.2285 | 0.1595 | 0.1498 |
| *Strongylodon macrobotrys* | 50-kb inversion clade | 0.2025 | 0.2126 | 0.2343 | 0.2058 | 0.1752 | 0.1724 |
| *Dipogon lignosus* | 50-kb inversion clade | 0.136 | 0.2157 | 0.2337 | 0.2087 | 0.1857 | 0.1621 |
| *Erythrina herbacea* | 50-kb inversion clade | 0.1407 | 0.2078 | 0.2379 | 0.1822 | 0.1613 | 0.1559 |
| *Dahlstedtia pinnata* | 50-kb inversion clade | 0.1927 | 0.2199 | 0.2299 | 0.2398 | 0.1622 | 0.17 |
| *Platycyamus regnellii* | 50-kb inversion clade | 0.2143 | 0.1933 | 0.2321 | 0.2199 | 0.177 | 0.1719 |
| *Canavalia villosa* | 50-kb inversion clade | 0.1757 | 0.1884 | 0.2351 | 0.2076 | 0.1479 | 0.1669 |
| *Amburana cearensis* | Other legumes | 0.097 | 0.1712 | 0.177 | 0.1392 | 0.1825 | 0.1582 |
| *Myrocarpus frondosus* | Other legumes | 0.045 | 0.1665 | 0.1703 | 0.135 | 0.1811 | 0.1705 |
| *Myrospermum sousanum* | Other legumes | 0.0738 | 0.1725 | 0.14 | 0.1554 | 0.1822 | 0.1686 |
| *Myroxylon balsamum* | Other legumes | 0.0709 | 0.1588 | 0.1678 | 0.1371 | 0.1807 | 0.15 |
| *Dipteryx alata* | Other legumes | 0.1003 | 0.145 | 0.1476 | 0.1531 | 0.1712 | 0.19 |
| *Ateleia glazioveana* | Other legumes | 0.1035 | 0.1691 | 0.1643 | 0.1369 | 0.1664 | 0.1593 |

**Table S6.** Genbank accession and voucher numbers for included taxa.

| Taxon Name | Clade Name | Voucher | SRA Accession Number | Mitogenome CDS Accession Numbers |
|---|---|---|---|---|
| *Cercis canadensis* | Cercidoideae | OneKP | ERR706845 | Accession numbers will be provided during review |
| *Parkinsonia aculeata* | Caesalpinioideae | I.S. Choi 1906007 (TEX-LL) | SAMN41673304 | “” |
| *Andira humilis* | Andira | D.Cardoso 4756 (HUEFS) | SAMN41673274 | “” |
| *Hymenolobium janeirense* | Andira | D.Cardoso 4791 (HUEFS) | SAMN41673297 | “” |
| *Exostyles venusta* | Exostyleae | D.Cardoso 3800 (RB) | SAMN41673292 | “” |
| *Harleyodendron unifoliolatum* | Exostyleae | L.P. de Queiroz 15470 (HUEFS) | SAMN41673294 | “” |
| *Holocalyx balansae* | Exostyleae | J.E.Meireles s.n. (RB no. 177569) | SAMN41673296 | “” |
| *Zollernia ilicifolia* | Exostyleae | M.J.Falcão 93 (RB) | SAMN41673317 | “” |
| *Luetzelburgia bahiensis* | Vataireoid | D.Cardoso 4787 (HUEFS) | SAMN41673298 | “” |
| *Sweetia fruticosa* | Vataireoid | D.Cardoso 2222 (RB) | SAMN41673313 | “” |
| *Vatairea guianensis* | Vataireoid | D.Cardoso 2195 (RB) | SAMN41673316 | “” |
| *Dermatophyllum secundiflorum* | Genistoid s.l. | I.S.Choi 1906003 (TEX-LL) | SAMN41673287 | “” |
| *Bowdichia virgilioides* | Genistoid s.l./Leptolobieae | L.F.G.Silva 30 (RB) | SAMN41673277 | “” |
| *Diplotropis ferruginea* | Genistoid s.l.//Leptolobieae | D.Cardoso 4739 (HUEFS) | SAMN41673288 | “” |
| *Camoensia scandens* | Genistoid s.l.//Camoensieae | D.Cardoso 2356 (HUEFS) | SAMN41673278 | “” |
| *Baptisia leucophaea* | Genistoid s.l.//Sophoreae | R.K.Jansen and T.A.Ruhlman 9 (TEX-LL) | SAMN41673276 | “” |
| *Cyclolobium brasiliense* | Genistoid s.l.//Brongniartieae | H.C. de Lima 5692 (RB) | SAMN41673283 | “” |
| *Harpalyce brasiliana* | Genistoid s.l.//Brongniartieae | D.Cardoso 4746 (HUEFS) | SAMN41673295 | “” |
| *Poecilanthe parviflora* | Genistoid s.l.//Brongniartieae | H.C. de Lima 2816 (RB) | SAMN41673309 | “” |
| *Tabaroa caatingicola* | Genistoid s.l.//Brongniartieae | D.Cardoso 4785 (HUEFS) | SAMN41673314 | “” |
| *Ormosia bahiensis* | Genistoid s.l.//Ormosieae | D.Cardoso 4790 (HUEFS) | SAMN41673303 | “” |
| *Acosmium lentiscifolium* | Dalbergioid s.l. | D.Cardoso 2216 (RB) | SAMN41673272 | “” |
| *Centrolobium tomentosum* | Dalbergioid s.l. | L.F.G.Silva 56 (RB) | SAMN41673280 | “” |
| *Dalbergia nigra* | Dalbergioid s.l. | J.E.Meireles s.n. (RB no. 322876) | SAMN41673285 | “” |
| *Platymiscium pubescens* | Dalbergioid s.l. | L.F.G.Silva 24 (RB) | SAMN41673308 | “” |

| | | | | |
|---|---|---|---|---|
| *Poiretia bahiana* | Dalbergioid s.l. | D.Cardoso 4795 (HUEFS) | SAMN41673310 | “” |
| *Stylosanthes scabra* | Dalbergioid s.l. | A.Oliveira 18 (HUEFS) | SAMN41673312 | “” |
| *Zornia myriadena* | Dalbergioid s.l. | N.P.Smith 45 (HUEFS) | SAMN41673318 | “” |
| *Dalea aurea* | Amorpheae | R.K.Jansen and T.A.Ruhlman 29 (TEX-LL) | SAMN41673286 | “” |
| *Eysenhardia texana* | Amorpheae | R.K.Jansen and T.A.Ruhlman 15 (TEX-LL) | SAMN41673293 | “” |
| *Alhagi sparsifolia* | IRLC/Hedysaroid | N/A | SRR1607745 | “” |
| *Trifolium repens* | IRLC/Vicioid | N/A | ERR351507 | CP125852.1 |
| *Coursetia rostrata* | NPAAA/Robinioid | D.Cardoso 4783 (HUEFS) | SAMN41673282 | “” |
| *Clitoria mariana* | NPAAA/Millettioid/Clitoriinae | R.K.Jansen and T.A.Ruhlman 27 (TEX-LL) | SAMN41673281 | “” |
| *Periandra mediterranea* | NPAAA/Millettioid/Clitoriinae | D.Cardoso 4738 (HUEFS) | SAMN41673305 | “” |
| *Macroptilium erythroloma* | NPAAA/Millettioid/Phaseolinae | D.Cardoso 4743 (HUEFS) | SAMN41673299 | “” |
| *Phaseolus acutifolius* | NPAAA/Millettioid/Phaseolinae | R.K.Jansen and T.A.Ruhlman 146 (TEX-LL) | SAMN41673306 | “” |
| *Strongylodon macrobotrys* | NPAAA/Millettioid/Phaseolinae | D.Cardoso 4656 (RB) | SAMN41673311 | “” |
| *Dipogon lignosus* | NPAAA/Millettioid | R.K.Jansen and T.A.Ruhlman 484 (TEX-LL) | SAMN41673289 | “” |
| *Erythrina herbacea* | NPAAA/Millettioid | I.S.Choi 1906006 (TEX-LL) | SAMN41673291 | “” |
| *Dahlstedtia pinnata* | NPAAA/Millettioid | J.R.Mattos 622 (RB) | SAMN41673284 | “” |
| *Platycyamus regnellii* | NPAAA/Millettioid | L.F.G.Silva 87 (RB) | SAMN41673307 | “” |
| *Canavalia villosa* | NPAAA/Millettioid /Diocleae | R.K.Jansen and T.A.Ruhlman 143 (TEX-LL) | SAMN41673279 | “” |
| *Amburana cearensis* | ADA/Amburaneae | N.L.Nunes 19 (RB) | SAMN41673273 | “” |
| *Myrocarpus frondosus* | ADA/Amburaneae | D.Cardoso 2204 (RB) | SAMN41673300 | “” |
| *Myrospermum sousanum* | ADA/Amburaneae | I.S.Choi 1906001 (TEX-LL) | SAMN41673301 | “” |
| *Myroxylon balsamum* | ADA/Amburaneae | D.Cardoso 2207 (RB) | SAMN41673302 | “” |
| *Dipteryx alata* | ADA/Dipterygeae | Equipe Arboreto s.n. (RB no. 435018) | SAMN41673290 | “” |
| *Ateleia glazioveana* | Swartzieae | H.C. de Lima 6279 (RB) | SAMN41673275 | “” |
| *Trischidium mole* | Swartzieae | D.Cardoso 4778 (HUEFS) | SAMN41673315 | “” |

**Table S7.** Nuclear-encoded control genes in glycolysis (N-gly), cell cycle (N-cc), cytosolic ribosomes (N-cr) gene sets and random nuclear proteins (N-rand) chosen from EnsemblPlants (Yates et al. 2022).

| Gene group | Gene symbol | Gene description | Uniprot ID |
|---|---|---|---|
| N-gly | G3PC2 | Glyceraldehyde-3-phosphate dehydrogenase GAPC2, cytosolic | Q9FX54 |
| N-gly | PGI1 | Glucose-6-phosphate isomerase | A0A090MHY5 |
| N-gly | PFKA3 | ATP-dependent-6-phosphofructokinase 3 | Q94AA4 |
| N-gly | ALFC6 | Fructose-bisphosphate aldolase 6, cytosolic | Q9SJQ9 |
| N-gly | TPIS | Triosephosphate isomerase, cytosolic | P48491 |
| N-gly | PFK | ATP-dependent-6-phoshofructokinase | A0A178V315 |
| N-gly | G6PI | Glucose-6-phosphate isomerase, cytosolic | P34795 |
| N-gly | PMG1 | 2,3-bisphosphoglycerate-independent phosphoglycerate mutase 1 | O04499 |
| N-gly | HKL1 | Phosphotransferase | A0A1P8ASN0 |
| N-gly | HXK2 | Hexokinase-2 | P93834 |
| N-cc | RBR1 | Retinoblastoma-related protein 1 | Q9LKZ3 |
| N-cc | CKS1 | Cyclin-dependent kinases regulatory subunit 1 | O23249 |
| N-cc | CDC2C | Cyclin-dependent kinase C-2 C | B5X564 |
| N-cc | TPX2 | Protein TPX2 | F4I2H7 |
| N-cc | DPA | Transcription factor-like protein DPA | Q9FNY3 |
| N-cc | KRP1 | Cyclin-dependent kinase inhibitor 1 | Q67Y93 |
| N-cr | RPL10AA | 60S ribosomal protein L10a-1 | Q8VZB9 |
| N-cr | RPL12A | 60S ribosomal protein L12-1 | P50883 |
| N-cr | RPL17A | 60S ribosomal protein L17-1 | Q93VI3 |
| N-cr | RPL23AA | 60S ribosomal protein L23a-1 | Q8LD46 |
| N-cr | RPL26B | 60S ribosomal protein L26-2 | Q9FJX2 |
| N-cr | RPL3A | 60S ribosomal protein L3-1 | P17094 |
| N-cr | RPL3B | 60S ribosomal protein L3-2 | P22738 |
| N-cr | RPL4A | 60S ribosomal protein L4-1 | Q9SF40 |
| N-cr | RPL7A | 60S ribosomal protein L7-1 | Q9SAI5 |
| N-cr | RPL8B | 60S ribosomal protein L8-2 | Q4PSL7 |
| N-cr | RPP0C | 60S acidic ribosomal protein P0-3 | P57691 |
| N-cr | RPS13B | 40S ribosomal protein S13-2 | P59224 |
| N-cr | RPS15AE | 40S ribosomal protein S15a-5 | Q9M0E0 |
| N-cr | RPS20C | 40S ribosomal protein S20-1 | P49200 |
| N-cr | RPS5B | 40S ribosomal protein S5-2 | P51427 |
| N-cr | RPS9C | 40S ribosomal protein S9-2 | Q9FLF0 |
| N-cr | RPSaA | 40S ribosomal protein Sa-1 | Q08682 |
| N-rand | MDN1 | Midasin | A0A1P8AUY4 |
| N-rand | REV1 | DNA repair protein REV1 | A3EWL3 |

| | | | |
|---|---|---|---|
| N-rand | 1MMP | Metalloendoproteinase 1-MMP | O23507 |
| N-rand | SRS2 | ATP-dependent DNA-helicase SRS2 | D1KF50 |
| N-rand | IDM1 | Increased DNA methylation 1 | F4IXE7 |
| N-rand | CERK1 | Chitin elicitor receptor kinase 1 | A8R7E6 |
| N-rand | LNK1 | Protein LNK1 | A8MQN2 |
| N-rand | RPP1 | Probable disease resistance protein RPP1 | F4J339 |
| N-rand | SAM2 | S-adenosylmethionine synthase 2 | P17562 |
| N-rand | BCB | Blue copper protein | Q07488 |
| N-rand | BZIP34 | Basic leucine zipper 34 | F4IN23 |
| N-rand | TK1B | Thymidine kinase b | F4KBF5 |
| N-rand | SAHH1 | Adenosylhomocysteinase 1 | O23255 |
| N-rand | CYP98A3 | Cytochrome P450 98A3 | O22203 |
| N-rand | KNL2 | Kinetochore-associated protein KNL-2 homolog | F4KCE9 |
| N-rand | AGL15 | Agamous-like MADS-box protein AGL15 | Q38847 |
| N-rand | GL2 | Homeobox-leucine zipper protein GLABRA 2 | P46607 |
| N-rand | APY6 | Probable apyrase 6 | O80612 |
| N-rand | HUB1 | E3 ubiquitin-protein ligase BRE1-like 1 | Q8RXD6 |
| N-rand | GRF1 | 14-3-3-like protein GF14 chi | P42643 |

**Table S8.** Summary of datasets compiled for analysis. 17 plastid genes were the plastid-encoded ribosomal genes (CpRP) used in Tressel et al. (2025).

| Dataset | Number of genes | Genome | OXPHOS Complexes |
|---|---|---|---|
| mt | 17 | mitochondrial | I,III,IV,V |
| mt-CI | 9 | mitochondrial | I |
| mt-CIII | 1 | mitochondrial | III |
| mt-CIV | 2 | mitochondrial | IV |
| mt-CV | 5 | mitochondrial | V |
| N-mt | 60 | nuclear | I,II,III,IV,V |
| N-mt-CI | 29 | nuclear | I |
| N-mt-CII | 7 | nuclear | II |
| N-mt-CIII | 9 | nuclear | III |
| N-mt-CIV | 6 | nuclear | IV |
| N-mt-CV | 9 | nuclear | V |
| N-mt contact | 42 | nuclear | I, III, IV, V |
| N-mt contact | 28 | nuclear | I, III, IV, V |
| Glycolysis | 10 | nuclear | none |
| Cell cycle | 6 | nuclear | none |
| Cytosolic-ribosomal | 17 | nuclear | none |
| Plastid | 17 | plastid | none |
